# Mucus transcriptional profiling as a minimally invasive approach to identify thermal stress in a stenothermal salmonid

**DOI:** 10.64898/2026.04.23.720280

**Authors:** A. Lazaro-Côté, T. C. Durhack, B. C. Kissinger, N. J. Mochnacz, K. M. Jeffries

**Affiliations:** Department of Biological Sciences, University of Manitoba, Winnipeg, Manitoba, Canada; Fisheries and Oceans Canada, Freshwater Institute, Winnipeg, Manitoba, Canada; Department of Biological Sciences, University of Calgary, Calgary, Alberta, Canada; fRI Research, Water and Fish Program, Hinton, Alberta, Canada

**Author notes:** **Corresponding author:** Analisa Lazaro-Côté.

**Keywords:** high-throughput qPCR, machine learning, random forest, sex differences, *Salvelinus confluentus*, bull trout

## Abstract

Global climate change has increased the frequency and severity of stressful temperatures that freshwater fishes experience, necessitating rapid and sensitive methods to monitor wild populations. Tissues used to measure transcriptional responses traditionally involved invasive or lethal sampling, which may be undesirable for imperilled species. Epidermal mucus offers a non-lethal and minimally invasive alternative, but whether thermal thresholds can be detected in mucus to identify fish experiencing thermal stress is unclear. Bull trout (*Salvelinus confluentus*) are a legally protected salmonid and cold-water specialist, generally occupying waters 12 °C and below, with higher temperatures resulting in cellular stress. Therefore, we measured a suite of 56 genes using high-throughput qPCR to compare machine learning classifiers developed with transcriptional profiles of epidermal mucus, gill, liver, and muscle to classify laboratory reared juvenile bull trout as below (9 °C, 12 °C) or above (15 °C, 18 °C) cellular thermal thresholds. Mucus profiles most resembled gills but represented an intermediate transcriptional response to all tissues. A reduced biomarker panel of 10 genes in mucus assigned fish to stress categories with 94.1% (95% CI = 71.3–99.9%) accuracy, which was comparable to gill (100.0%, CI = 82.4– 100%), liver (95.0%, CI = 75.1–99.9%), and muscle (100.0%, CI = 80.5–100.0%). Sex-specific temperature effects were evident in all tissues, but less pronounced in mucus and gill than in liver and muscle. Our findings demonstrate that transcriptional profiling of mucus can reliably identify individuals experiencing thermal stress, highlighting the promise of this non-lethal approach for monitoring at-risk species.

## Introduction

Global climate change is increasing the frequency and severity of thermal stress experienced by ectotherms (Sunday et al. 2014; Alfonso et al. 2021), creating a need for rapid and sensitive tools to assess the physiological status of wild populations. The use of transcriptional profiling to evaluate physiological responses to ecological stressors is increasing, in part because changes in gene expression can indicate sublethal stress before organismal-level effects become apparent (Connon et al. 2018). Transcriptional profiling has traditionally relied on tissues obtained via dissections. However, there is growing interest in less invasive approaches, particularly to assess the status of imperilled species (Jeffries et al. 2021; Lazaro-Côté et al. 2025).

Epidermal mucus, which can be collected rapidly with reduced harm to the animal (Sanahuja et al. 2023), has emerged as an alternative matrix for evaluating stress. In fishes, mucus is found on the outermost layer of the epidermis, gills, and gut. Epidermal mucus is primarily secreted by goblet cells and composed largely of water and glycoproteins (mucins), acting as a physical and chemical barrier between fish and their aquatic environment (Shephard 1994; Brinchmann 2016). Mucus plays a role in osmoregulation, parental care, locomotion, and chemical communication in addition to protecting against abrasions, pathogens, and pollutants (Shephard 1994; Brinchmann 2016). Environmental stressors, including elevated temperatures and low dissolved oxygen (Weinrauch et al. 2025), pathogens (Ren et al. 2015), and contaminants (Greer et al. 2019; Andrzejczyk et al. 2022), alter the mRNA transcript levels in mucus which enables an assessment of the physiological state of the animal. However, it is unclear whether finer scale thermal stress thresholds are detectable in mucus and the predictive accuracy of mucus transcriptional profiling to identify stress in fish has not been examined relative to other tissues.

When paired with machine learning approaches, such as random forest classifiers that can be used to assign individuals to stress categories based on a panel of biomarkers, transcriptional profiling can identify stressed individuals with high accuracy (von Biela et al. 2023; Akbarzadeh et al. 2024; Frommel et al. 2025). In Chinook salmon (*Oncorhynchus tshawytscha*) and coho salmon (*Oncorhynchus kisutch*), transcriptional profiles from white muscle biopsies were used to classify individuals experiencing thermal stress (von Biela et al. 2023). Similarly, gill transcriptional responses in coho salmon identified fish experiencing salinity, hypoxia, or thermal stress (Akbarzadeh et al. 2024), while profiles in chum salmon (*Oncorhynchus keta*) identified exposure to acidification or nutritional stress (Frommel et al. 2025). Across studies, random forest classifiers achieved high prediction accuracy with as few as 4 to 13 gene biomarkers, demonstrating the potential of high-throughput transcriptional profiling to support ecological monitoring. Yet, this machine learning approach has not been used with mucus to facilitate interpretation of transcriptional biomarkers in fish.

Developmental stage, sex, and thermal history can further influence physiological responses to rising temperatures (Geffroy and Wedekind 2020, Mayer et al. 2023). Although sex-specific differences in mucus composition have been documented (Wen et al. 2020; Roosta et al. 2023; Valero et al. 2024), it is not known how mucus transcriptional responses to thermal stress differ between sexes. Sex-specific mortality associated with thermal stress has been thoroughly studied in adult Pacific salmon, with mounting evidence that females often experience greater mortality rates (Little et al. 2020; Hinch et al. 2021). Mature females are believed to be more vulnerable to stress due to energy exhaustion during gamete formation, impaired cardiorespiratory function, slower recovery from stress, and immunosuppression (Hinch et al. 2021). For instance, exposure to elevated temperatures prior to migration resulted in greater mortality for female compared to male sockeye salmon (*Oncorhynchus nerka)* migrating to spawning grounds (Crossin et al. 2008). Less is known about whether juveniles experience sex-biased mortality due to thermal stress. Despite these patterns, sex is often not reported nor accounted for in thermal acclimation studies (reviewed in Pottier et al. 2021), limiting our understanding of sex-specific transcriptional thermal thresholds.

Bull trout (*Salvelinus confluentus*) are cold-water specialists whose abundance and distribution are increasingly threatened by climate warming, interspecific competition with other salmonids, and habitat degradation (Fisheries and Oceans Canada 2020). Their protected status throughout most of North America necessitates minimally invasive methods to assess the physiological status of wild populations. Transcriptional profiling of gill, liver, and muscle in temperature-acclimated young-of-year bull trout has identified cellular thermal thresholds between 12 °C and 15 °C (Best et al. 2025). However, we currently do not know if similar transcriptional thresholds are reflected in epidermal mucus or if sex may influence these thresholds. Therefore, we tested the hypotheses that (1) transcriptional profiling of epidermal mucus can be used to detect thermal thresholds and accurately classify fish experiencing sublethal thermal stress, and (2) females experience cellular stress at different temperatures than males. We predicted that mucus transcriptional profiles would (i) most resemble those in gill, with a thermal threshold between 12 °C and 15 °C, as both tissues interface directly with the aquatic environment, (ii) classify thermally stressed fish with accuracy comparable to gill, liver, and muscle, and (iii) reveal lower transcriptional thermal thresholds in females than in males. Developing sensitive, non-lethal approaches to detect sublethal stress in fish will enable early intervention before adverse effects are observed at the organismal or population level, supporting conservation and management initiatives.

## Methods

### Fish Holding

All animal handling was approved by the Fisheries and Oceans Canada Ontario, Prairie, and Arctic Animal Care Committee (Animal use protocol #OPA-ACC-2024-21). Bull Trout were reared from gametes collected in September 2021 from a wild, montane population in Smith-Dorrien Creek, Alberta, Canada (for more details see Kissinger et al. 2025). In April 2024, fish (aged 2.25 years) were randomly assigned to one of six 200-L tanks in groups of 45 and acclimated to either 9 °C, 12 °C, 15 °C, or 18 °C for 4 weeks. The start date for each temperature acclimation was staggered over four weeks to facilitate sampling. Fish were fed a daily ration of 1% total tank mass (#2 pellet EWOS Pacific: Complete Fish Feed for Salmonids, Cargill). Throughout rearing and experiments, overhead lighting was set to a 12h light: 12h dark cycle (lights on 06:00-18:00) using adjustable lights to simulate the colour and intensity of sunrise and sunset.

Fish were fasted for 24 h prior to sampling. For each acclimation temperature, bull trout (n = 15/temperature) were randomly selected and euthanized in 300 mg/L MS-222 buffered with 600 mg/L sodium bicarbonate. Epidermal mucus samples were collected from each fish using a PurFlock Ultra Sterile Flocked Collection Device (Puritan Medical Products) to swab from lateral line to dorsal fin across their body length. Fork length (mm) and mass (g) were recorded, condition factor was calculated (K=mass/(length^3^) x 100), then each fish was dissected for gill filaments from the second arch, liver, and epaxial white muscle, during which time sex was determined by visual inspection of gonads. Whole swab tips and tissue samples were placed in RNA*later* (Invitrogen) and stored at -80 °C.

To determine if mucus sampling increased mortality risk, 20 fish were anaesthetized in 80 mg/L MS-222 buffered with 160 mg/L sodium bicarbonate. Swabs of epidermal mucus were collected from 10 randomly selected fish, leaving the other 10 fish with their mucus layer undisturbed. Before returning fish to tanks to recover, each fish was tagged with one of two colours of visible implant elastomers (Northwest Marine technology, Inc.) near the base of the dorsal fin to identify which fish were sampled for mucus and which ones were not. Survival was monitored for 96 h, and sex was determined by visually inspecting gonads when fish were later euthanized.

### Total RNA extraction, cDNA synthesis, and mRNA transcript abundance

Total RNA was successfully extracted from 30–40 mg of gill (n = 60), liver (n = 60), muscle (n = 52), and mucus swabs (n = 51) using the MagMAX™ *mir*Vana™ Total RNA Isolation Kit (Applied Biosystems) with the KingFisher Duo Prime Purification System (Thermo Scientific) following manufacturer’s instructions. Tissues were homogenized in 600 ul (gill, liver, mucus) or 200 ul (mucus swab) of lysis buffer at 50 hz for 5 min using a TissueLyser II (Qiagen). RNA quantity and quality was determined with a NanoDrop One (Thermo Scientific). We synthesized cDNA (High Capacity cDNA Reverse Transcription Kit, Applied Biosystems) with 1000 ng (gill, liver) or 350 ng (muscle, mucus) of RNA treated with ezDNase (Invitrogen).

Custom Taqman OpenArray Real-Time PCR plates (Thermo Scientific) developed for salmonids were used to quantify the mRNA transcript abundance of 56 genes related to stress responses, detoxification, growth and metabolism, osmoregulation, apoptosis, and immune function (Islam et al. 2026; Table S1). Briefly, 2.5 ul of 5X-diluted cDNA was mixed with 2.5 ul TaqMan™ OpenArray™ Real-Time PCR Master Mix then loaded onto an OpenArray plate using the QuantStudio 12K Flex OpenArray AccuFill System (Applied Biosystems). The QuantStudio 12K Flex Real-Time PCR System (Applied Biosystems) was used to quantify mRNA transcript abundance in OpenArray plates.

Amplification score and Cq confidence were used to evaluate the quality of PCR amplification in each well. Target genes were removed from final analyses if over 50% of the wells were undetermined, had an amplification score < 1.24, or a Cq confidence < 0.8. For each tissue type, LinRegPCR (v.2021.2) was used to estimate primer efficiency for target genes using the baseline-corrected fluorescence values derived from amplification curves (Ruijter et al. 2009; Tuomi et al. 2010). Gene targets with a primer efficiency under 1.6 (80%) or over 2.4 (120%) were removed from analyses (Table S1). Following the 2^-ΔCT^ approach (Schmittgen and Livak 2008), transcript abundance in mucus, gill, and liver were normalized against the geometric mean of *rpl13a*, *rpl7*, *rps9*, whereas the geometric mean of *rpl7* and *rps9* was used for muscle. We omitted *ef1a* as a reference gene as its transcript abundance was not stable across treatments.

### Statistical analyses

All statistical analyses were conducted in R (v.4.3.2). Fork length, mass, and condition factor were analyzed with linear models (Gaussian distribution) using temperature, sex, and their interaction as categorical fixed variables. Normality and homoscedasticity of the residuals were assessed graphically. Post hoc analyses of main effects were performed with Tukey’s tests. Marginal (least squares) means with 95% confidence intervals are presented graphically.

To visualize tissue-specific differences in transcriptional profiles and determine the proximity of mucus transcriptional profiles to gill, liver, and muscle, a principal component analysis (PCA) was implemented using only genes that amplified in the mucus. Then, separate PCA’s were applied to each tissue using all genes that successfully amplified to visualize separation between fish acclimated to 9, 12, 15, or 18 °C. PCA’s were implemented using the package *FactoMineR* (Lê et al. 2008).

The onset of cellular thermal stress in bull trout is reportedly between 12 and 15 °C (Best et al. 2025). Therefore, bull trout acclimated to 9 or 12 °C were assigned as “below”, while bull trout acclimated to 15 or 18 °C were assigned as “above” the cellular stress threshold. For each tissue type, random forest classifiers were built with transcriptional profiles using the *randomForest* package (Liaw and Wiener 2002) to categorize fish as above or below the thermal stress threshold. We used random forest over other classification methods as it uses an ensemble of decision trees that use binary splits on predictor variables with accuracy often higher than other approaches (Fernández-Delgado et al. 2014). Transcriptional data were randomly split into 2/3 for training and 1/3 for testing classifiers using the *randomForest* function with a classification threshold set to 0.5, *mtry* = 2, and *ntree* = 1000. Increasing the number of trees beyond 1000 did not considerably reduce out-of-bag error rates.

The variable selection method implemented in the *VSURF* package (Genuer et al. 2015) was used to rank biomarkers and select the top 10 most important genes for classification (Figures S1–S4), because the permutation-based variable importance ranking of VSURF results in lower out-of-bag error rates with manageable computational costs relative to other procedures for datasets with binary outcomes (Speiser et al. 2019). For each tissue, the performance of one classifier using all target genes and one classifier using a reduced biomarker panel of the top 10 genes was assessed using the test data. We selected 10 genes because previous studies have reported high classification accuracy using fewer than 13 gene biomarkers (von Biela et al. 2023; Akbarzadeh et al. 2024; Frommel et al. 2025). The specificity (true negative rate), sensitivity (true positive rate), and accuracy (percentage of cases correctly classified) of each classifier was calculated using the *confusionMatrix* function from the *caret* package (Kuhn 2008). Additionally, a receiver operation characteristics (ROC) analysis was implemented using the function *roc* from the *pROC* package to determine the area under the curve (AUC) of each classifier (v.1.0-11; Robin et al. 2011).

To assess the influence of acclimation temperature and sex on tissue-specific mRNA transcript abundance, Bayesian generalized linear mixed models were implemented in the *MCMC.qpcr* (v1.2.4) package (Matz et al. 2013). The function *mcmc.qpcr* was used to fit Bayesian models because low-abundance genes are handled with a Poisson-lognormal error distribution, adding reference genes as priors narrows credible intervals without significantly shifting posterior means, and posterior probabilities for all pairwise comparisons are returned (Matz et al. 2013). Each model included gene, and gene-specific effects of temperature, sex, and their interaction as fixed effects, while variation among biological [sample] and technical replicates [gene:sample] was accounted for as random effects.

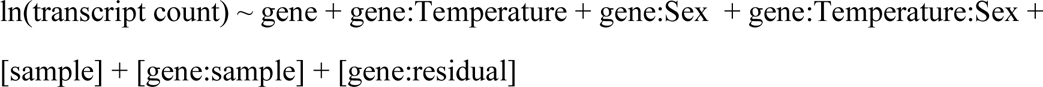

Mucus, gill, liver, and muscle transcript abundance were analyzed separately due to tissue-specific differences in primer efficiencies. Raw C_rt_ values were efficiency-corrected and converted to transcript counts using the function *cq2counts*. A Markov Chain Monte Carlo (MCMC) chain was run to fit a Poisson-lognormal generalized linear mixed model to the counts data. Models were fitted with MCMC settings of 13,000 iterations, with the first 3,000 iterations discarded, and a thinning interval of 10. This resulted in a target effective sample size of 1000 with the majority of parameters meeting this threshold. The stability of the reference genes and absence of global effects across acclimation temperatures and sex were determined visually by fitting a naïve model without priors. Models were then informed by incorporating reference genes as Bayesian priors, with a stability parameter set to a 1.2-fold change. The function *diagnostic.mcmc* was used to generate predicted vs observed plots, scale-location plots, and normal quantile-quantile plots to assess linearity, homoscedasticity, and normality of the residuals, respectively. For each gene, posterior means and 95% credible intervals for log_2_-transformed transcript abundance were plotted, and pairwise differences between acclimation temperatures and sexes were determined using the *HPDsummary* function.

## Results

### Body size and survival

Overall, condition factor was 8% higher for female bull trout compared to males (F_(1,52)_ = 6.90, *p* = 0.011), because females weighed 15% more (F_(1,52)_ = 5.31, *p* = 0.025) but did not significantly differ in length when compared to males (F_(1,52)_ = 1.85, *p* = 0.179). Temperature effects did not depend on sex as there was no significant interaction effect on fork length (F_(3,52)_ = 0.85, *p* = 0.474; Figure 1a), mass (F_(3,52)_ = 0.33, *p* = 0.802; Figure 1b), nor condition factor (F_(3,52)_ = 0.21, *p* = 0.890; Figure 1c) of bull trout. Condition factor differed significantly with temperature (F_(3,52)_ = 10.13, *p* < 0.001). Specifically, condition factor was significantly lower at 18 °C compared to 9 °C (t_(52)_ = 4.75, *p* = 0.001) and 12 °C (t_(52)_ = 3.71, *p* = 0.003), and lower at 15 °C compared to 9 °C (t_(52)_ = 3.79, p = 0.002). The decrease in condition factor with temperature was driven by a significant decrease in mass (F_(3,52)_ = 3.69, *p* = 0.017), rather than length (F_(3,52)_ = 0.43, *p* = 0.735), although a post-hoc test did not detect a significant decrease in mass between 9 °C and 18 °C (t_(52)_ = 2.57, *p* = 0.061). Survival was 100 % for bull trout males and females at each acclimation temperature regardless of whether mucus was sampled.

**Figure 1.**
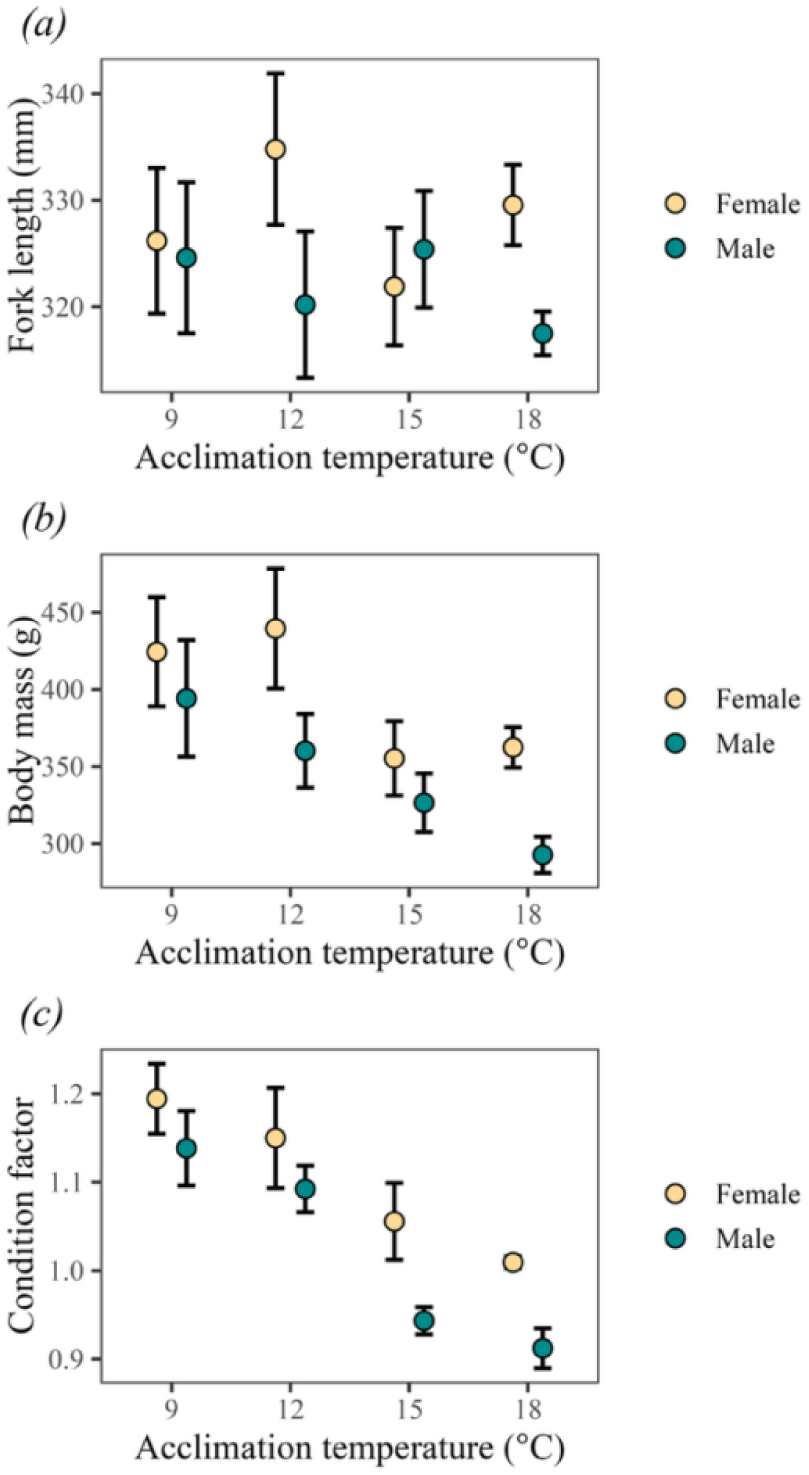
Mean and standard error of (a) fork length (mm), (b) body mass (g), and (c) condition factor of juvenile bull trout (*Salvelinus confluentus*) females (yellow circle; n = 9–10/temperature) and males (green circle; n = 5–6/temperature) acclimated to 9, 12, 15, or 18°C for four weeks.

### Random forest classification

Of the 52 gene targets (not including the four reference genes), 50 (96%) amplified in gill, 44 (85%) in liver, 42 (81%) in muscle, and 35 (67%) in mucus (Figures S5–S8). In the PCA, transcriptional profiles formed distinct tissue-specific clusters, with mucus positioned closest to gill on PC1 yet intermediate to all tissue types along PC1 and PC2 (Figure 2). For each tissue, transcriptional profiles were most similar for fish acclimated to 9 and 12 °C, and those acclimated to 15 and 18 °C (Figures 3a–d). Gill profiles from fish exposed to 15 °C and those held at 18 °C were further distinguishable along PC1 and PC2 (Figure 3b).

**Figure 2.**
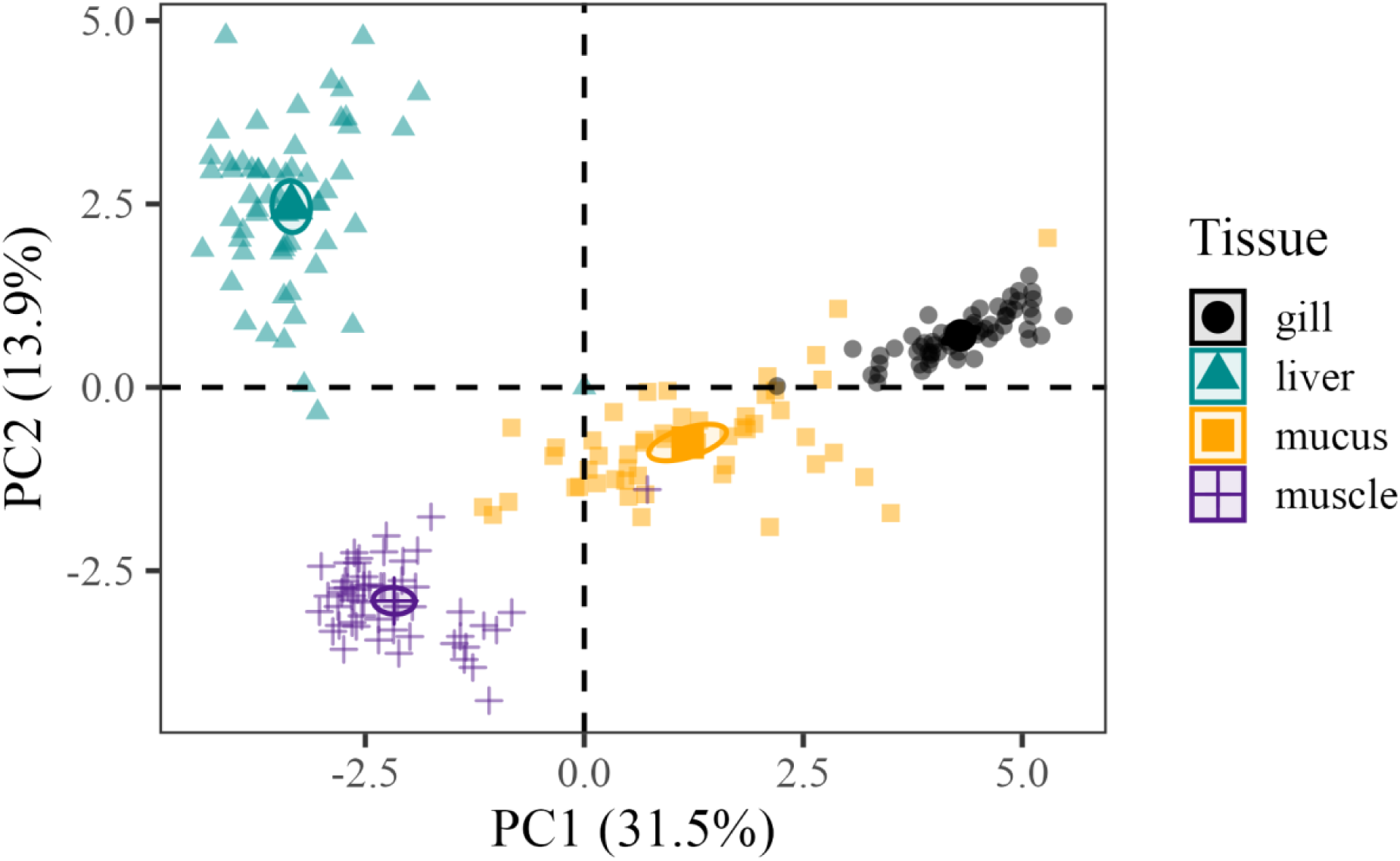
Principal component analysis of transcriptional responses in mucus, gill, liver, and muscle of juvenile bull trout (*Salvelinus confluentus*).

**Figure 3.**
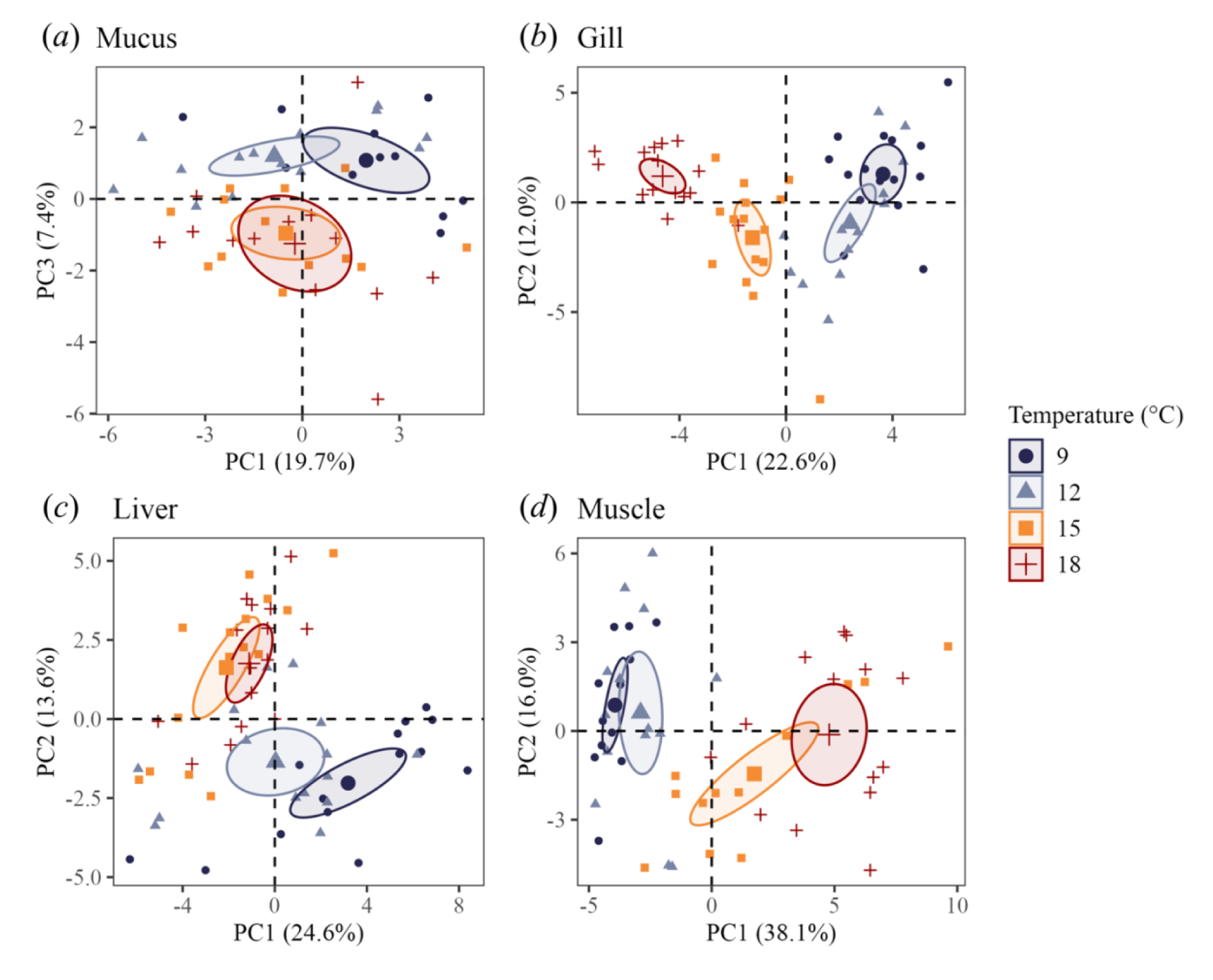
Principal component analysis of transcriptional responses in (a) mucus, (b) gill, (c) liver, and (d) muscle of juvenile bull trout (*Salvelinus confluentus*). Fish were exposed to temperatures between 9 °C and 18 °C for 4 weeks. The variance explained by each principal component (PC) is shown in brackets.

For each tissue, we selected the top 10 most informative genes for identifying juvenile bull trout experiencing thermal stress to generate reduced biomarker panels (Figures 4a, 5a, 6a, 7a). In epidermal mucus, biomarkers were associated with immune function (*il8*, *mhci*, *stat1*) and cellular stress responses (*chmp5ab*, *gstp1*, *hspa4*, *mta*, *mtb*), with additional genes involved in glycolysis (*pgk1*) and osmoregulation (*atp1b1*; Figure 4a). In gill, top biomarkers were primarily involved in stress responses (*casp3ab*, *cyp1a*, *hsp7c*, *mr*, *mta*), growth (*mmp2*, *mmp9*), metabolism (*aldoaa*, *cpt1a*), and immune function (*mhci*; Figure 5a). In liver, biomarkers were related to metabolism (*ampka*, *fasn*), growth (*ghr*, *igf1*, *igfbp1*, *mmp2*), and stress responses (*cyp1a*, *gstp1*, *mta*, *mtb*; Figure 6a). Finally, the most informative biomarkers in muscle were genes related to stress responses (*hsp70a*, *hsp7c*, *mtb*, *hspa4*, *cirbpa*, *cat*), immune function (*stat1*), protein degradation (*ctsd*), and apoptosis (*chmp5ab*; Figure 7a). Overall, metallothionein b (*mtb*), which helps protect against oxidative stress, emerged as an important biomarker in all tissues except gill. Mucus shared several other biomarkers, including *mta* and *gstp1* with liver, *mta* and *mhci* with gill, and *chmp5ab* and *hspa4* with muscle.

**Figure 4.**
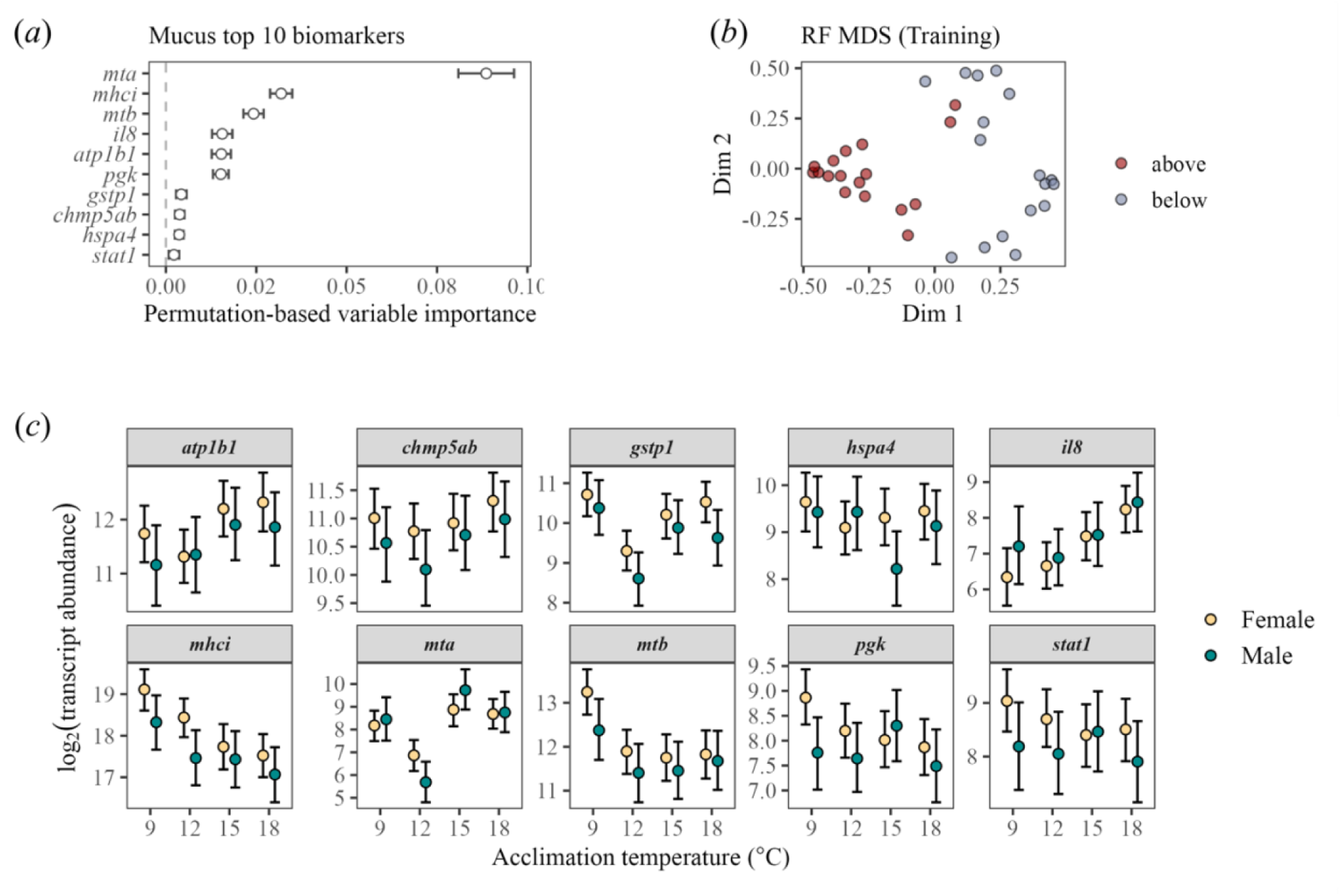
(a) Mucus top 10 genes ranked according to permutation-based variable importance. (b) Multi-dimensional scaling (MDS) plot of proximity matrix from training of random forest model to classify fish as above (red circle) or below (blue circle) their thermal threshold. (c) Mucus mRNA transcript abundance of top 10 genes of juvenile bull trout (*Salvelinus confluentus*) acclimated for four weeks to temperatures between 9 °C and 18 °C. Points represent posterior means and bars indicate 95% credible intervals for temperature effects in females (yellow circles; n = 9–10 females/temperature) and males (green circles; 5–6 males/temperature).

**Figure 5.**
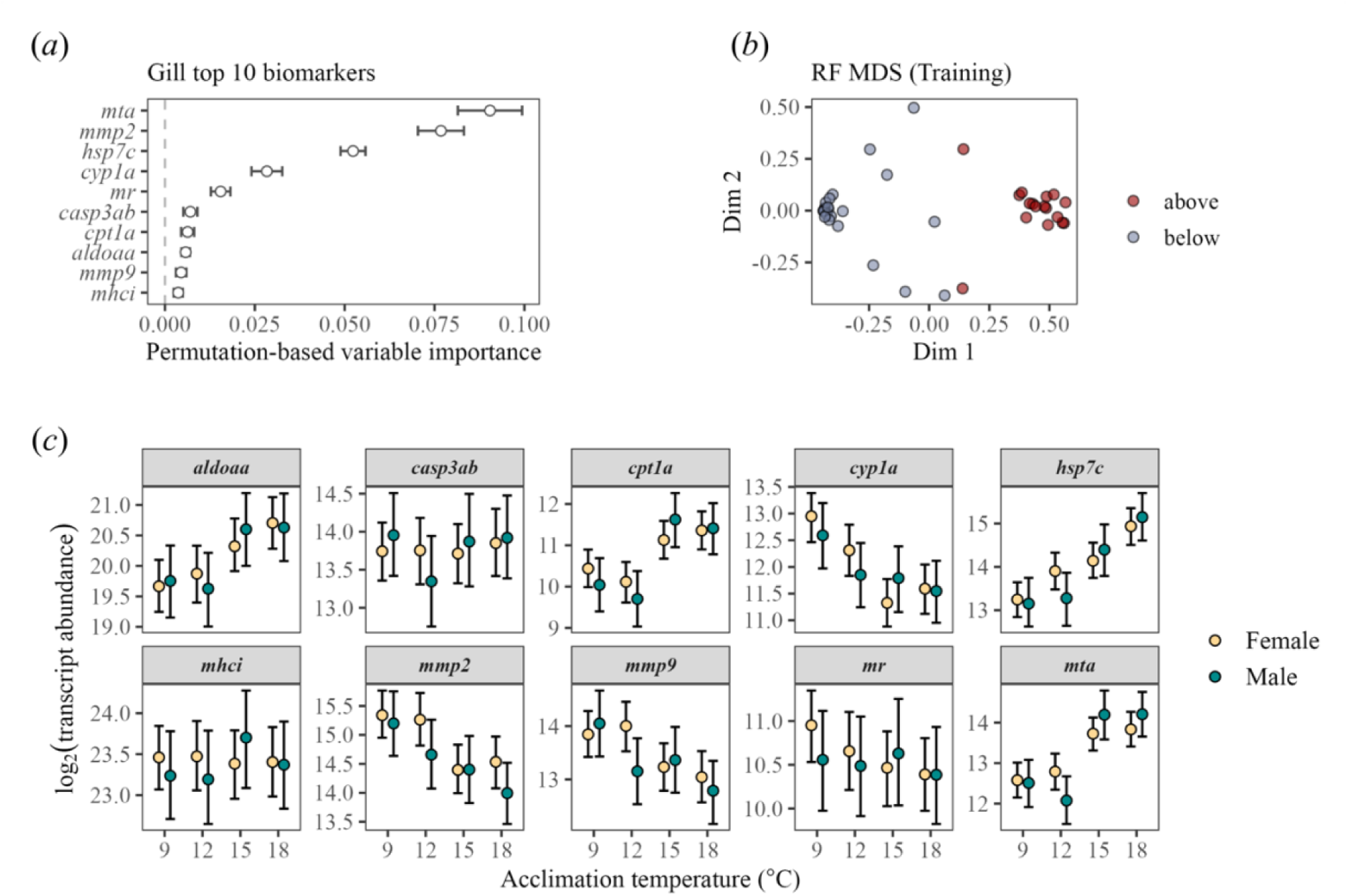
(a) Gill top 10 genes ranked according to permutation-based variable importance. (b) Multi-dimensional scaling (MDS) plot of proximity matrix from training of random forest model to classify fish as above (red circle) or below (blue circle) their thermal threshold. (c) Gill mRNA transcript abundance of top 10 genes of juvenile bull trout (*Salvelinus confluentus*) acclimated for four weeks to temperatures between 9 °C and 18 °C. Circles represent posterior means and bars show 95% credible intervals for temperature effects in females (yellow; n = 9–10 females/temperature) and males (green; 5–6 males/temperature).

**Figure 6.**
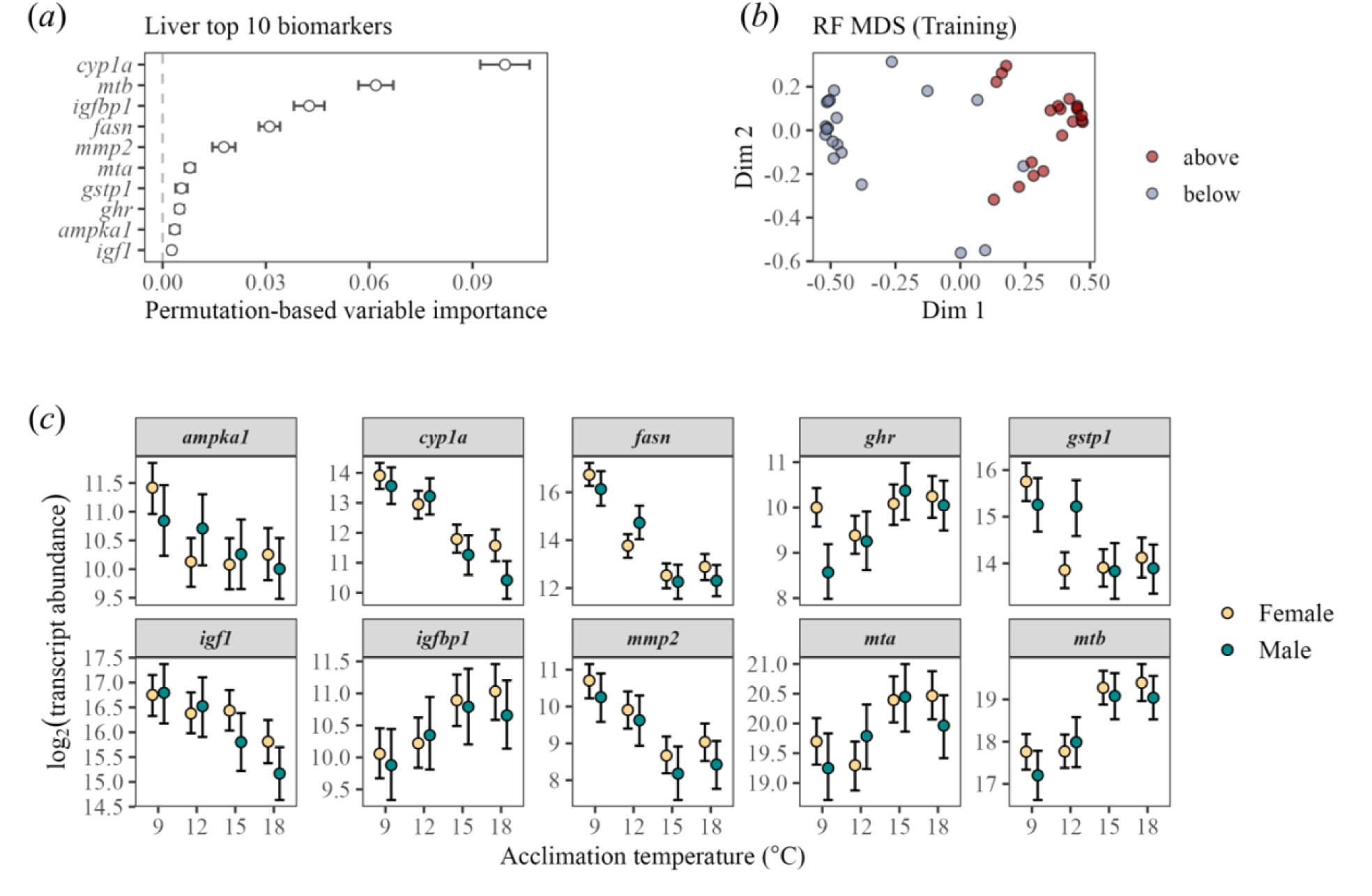
(a) Liver top 10 genes ranked according to permutation-based variable importance. (b) Multi-dimensional scaling (MDS) plot of proximity matrix from training of random forest model to classify fish as above (red circle) or below (blue circle) their thermal threshold. (c) Liver mRNA transcript abundance of top 10 genes of juvenile bull trout (*Salvelinus confluentus*) acclimated for four weeks to temperatures between 9 °C and 18 °C. Points represent posterior means and bars represent 95% credible intervals for temperature effects in females (yellow circles; 9–10 females/temperature) and males (green circles; 5–6 males/temperature).

**Figure 7.**
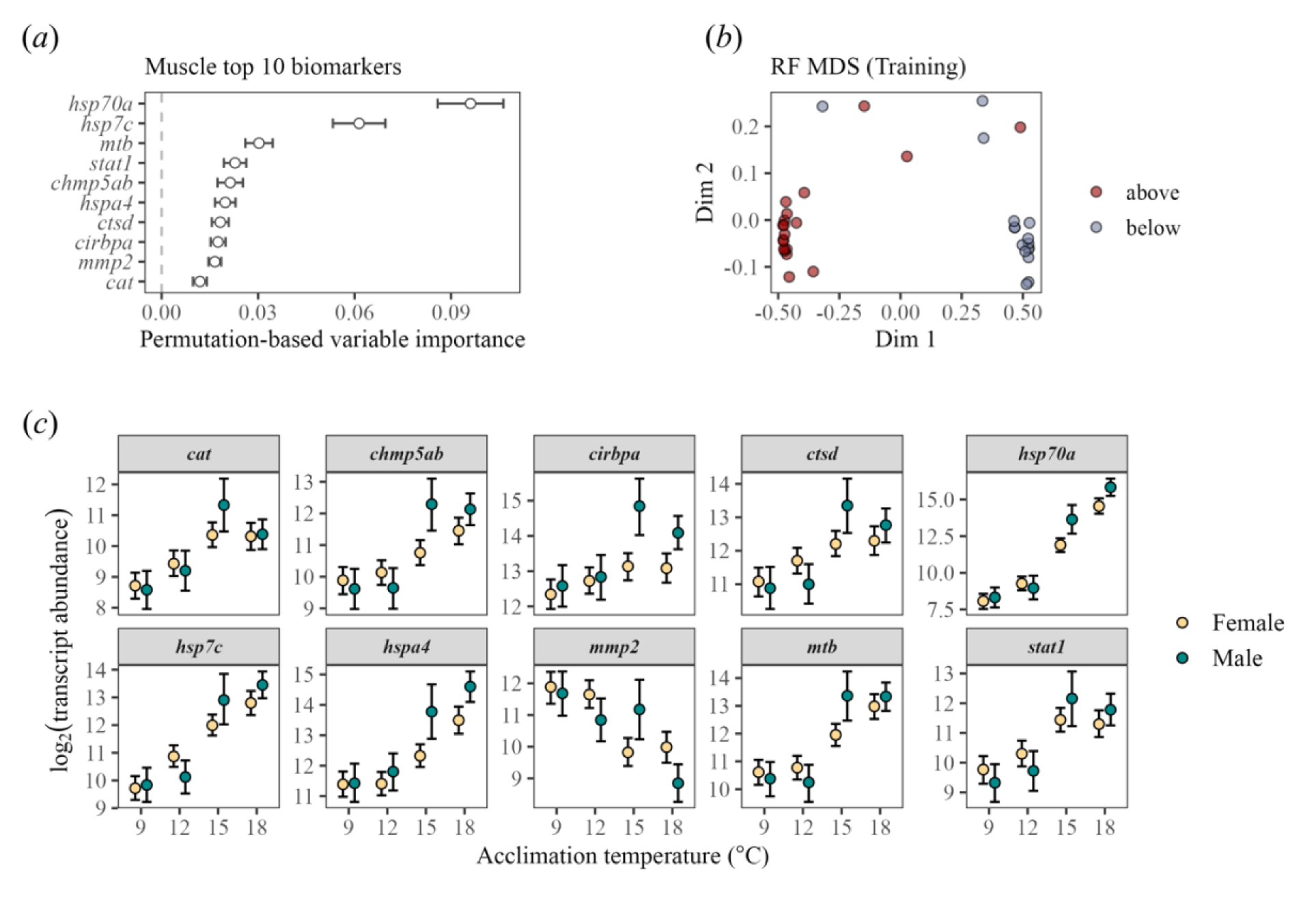
(a) Muscle top 10 genes ranked according to permutation-based variable importance. (b) Multi-dimensional scaling (MDS) plot of proximity matrix from training of random forest model to classify fish as above (red circle) or below (blue circle) their thermal threshold. (c) Muscle mRNA transcript abundance of top 10 genes of juvenile bull trout (*Salvelinus confluentus*) acclimated for four weeks to temperatures between 9 °C and 18 °C. Points represent posterior means and bars denote 95% credible intervals for temperature effects in females (yellow circles; n = 9–10 females/temperature) and males (green circles; 5–6 males/temperature).

For each tissue, the random forest classifier built using the reduced biomarker panel performed as well or better than the classifier using all genes when evaluated on the test dataset (Table 1). The area under the receiving operating characteristic curve (AUC) ranged from 0.97 to 1.00, indicating a high probability that the model will correctly classify fish according to thermal stress status (Table 1). The multidimensional scaling (MDS) plot of random forest proximities revealed clear separation between fish above or below their thermal threshold during model training, indicating strong discrimination by classifiers using reduced gene panels (Figures 4b, 5b, 6b, 7b). When categorizing fish based on mucus profiles, the classifier’s ability to correctly identify fish experiencing thermal stress (sensitivity = 88.9%) was lower than its ability to identify fish below their thermal threshold (specificity = 100.0%). However, mean classification accuracy using a reduced gene set in mucus (94.1%; CI = 71.3–99.9%) was between 0.9% and 5.9% lower than accuracies achieved with transcripts from gill (100.0%; CI = 82.4–100.0%), liver (95.0 %; CI = 75.1–99.9%), and muscle (100.0%; CI = 80.5–100.0%).

**Table 1.** Performance of random forest classifiers on test data using a threshold of 0.5. Performance is reported for classifiers using a full gene panel and a reduced panel of the top 10 genes for each tissue type.

| Tissue | Classifier | Sensitivity (%) | Specificity (%) | Accuracy (%) (95% CI) | AUC |
| --- | --- | --- | --- | --- | --- |
| Mucus | All genes (35) | 88.9 | 100.0 | 94.1 (71.3–99.9) | 98.6 |
|  | Top 10 genes | 88.9 | 100.0 | 94.1 (71.3–99.9) | 97.2 |
| Gill | All genes (50) | 92.3 | 100.0 | 94.7 (74.0–99.9) | 1.00 |
|  | Top 10 genes | 100.0 | 100.0 | 100.0 (82.4–100.0) | 1.00 |
| Liver | All genes (44) | 100.0 | 90.0 | 95.0 (75.1–99.9) | 0.98 |
|  | Top 10 genes | 100.0 | 90.0 | 95.0 (75.1–99.9) | 0.97 |
| Muscle | All genes (42) | 1.00 | 1.00 | 100.0 (80.5–100.0) | 1.00 |
|  | Top 10 genes | 1.00 | 1.00 | 100.0 (80.5–100.0) | 1.00 |

### Tissue-specific temperature and sex effects on transcript abundance

Overall, the interaction effect of acclimation temperature and sex on the transcript abundance of genes included in the reduced biomarker panels were tissue specific. In the mucus of both sexes, transcript abundance for markers of oxidative stress (*gstp1*, *mta*) was lowest at 12 °C compared to all other temperatures (Figure 4c). There were no within-temperature effects of sex on mucus transcriptional profiles (Figure S5). However, rising temperatures impacted immune-related genes (*il8*, *mhci*) and an additional oxidative stress marker (*mtb*) in females, but these effects were less pronounced in males. Specifically, increasing temperature was associated with the upregulation of *il8* and the downregulation of *mhci* and *mtb* transcripts. Evidence was weaker for temperature effects on *atp1b1*, *chmp5ab*, *hsp4a*, *pgk*, or *stat1* in mucus although they remained informative biomarkers to classify thermally stressed bull trout.

In gill, genes involved in fatty acid transport (*cpt1a*), protein repair (*hsp7c*), and oxidative stress (*mta*) were upregulated in both sexes when temperatures exceeded 15 °C (Figure 5c). Like mucus, there was no effect of sex within a given temperature on gill transcript abundance (Figure S6). However, the transcript abundance of an enzyme involved in glycolysis (*aldoaa*) increased, whereas those involved in xenobiotic metabolism (*cyp1a*) and tissue remodelling (*mmp2*) decreased in females when temperatures exceeded 12 °C. In contrast, temperature effects on *aldoaa* and *cyp1a* were weaker in males, and downregulation of *mmp2* was not observed until 18 °C. There was little support for temperature-related effects on genes associated with apoptosis (*casp3ab*), immune function (*mhci*), or cortisol signalling (*mr*) in gills, but they remained informative for accurate machine learning classification (Figure 4a).

Genes involved in growth signalling (*igf2*), xenobiotic metabolism (*cyp1a*), and tissue remodelling (*mmp2*) decreased, whereas markers of oxidative stress increased (*mta*, *mtb*) in the liver of both sexes when acclimation temperatures exceeded 15 °C (Figure 6c). In females, transcripts of a cellular energy sensor (*ampka1*) decreased and those related to growth regulation (*igfbp1*) increased with temperature, while evidence for temperature effects on these genes was weaker in males. Similarly, genes involved in fatty acid transport (*fasn*) and the antioxidant response (*gstp1*) were downregulated at 12 °C in females, but not until 15 °C in males. In contrast, the transcript abundance for growth hormone receptor (*ghr*) in females was greater at 9 °C and remained stable across temperatures but increased with temperature in males. Beyond the top biomarkers in liver, the response to acclimation temperature of several genes differed among sexes (Figure S7). The transcript abundance of genes involved in fatty acid transport (*cpt1a*) was consistently higher in females across all temperatures. In the liver of females relative to males, genes associated with growth signalling (*ghr*) and cortisol regulation (*hsd11b2*) were upregulated at 9 °C and 18 °C, respectively, whereas detoxification (*gstp1*) was downregulated at 12 °C (Figure S7).

Of the reduced biomarker panel for muscle, genes related to stress responses (*hsp70a*, *hsp7c*, *mtb*, *hspa4*, *cirbpa*, *cat*), immune function (*stat1*), protein degradation (*ctsd*), and apoptosis (*chmp5ab*) increased with acclimation temperature (Figure 7c). Conversely, the transcript abundance of a gene involved in tissue remodelling (*mmp2*) decreased with temperature. This temperature-dependent shift in transcriptional regulation occurred at 15 °C for most of the top biomarkers except for a temperature responsive RNA-binding protein (*cirbpa*) which did not vary considerably across temperatures for females. In contrast, transcripts involved in tissue remodelling (*mmp2*) decreased at 15 °C in females but not until 18 °C in males. Beyond the top biomarkers, transcript abundance was generally higher in males than females acclimated to elevated temperatures. Specifically, males upregulated genes associated with apoptosis (*chmp5ab*, *pdcd10ab*), energy substrate metabolism (*ldhb*, *lpl*, *ampka*), detoxification (*mtb*, *nfe2l2*), osmoregulation (*atp1b1*), signal transduction (*cam*, *ghr*, *hsd11b2*), and cellular stress (*hsp70a*, *hspa4, cirbpa*) when temperatures exceeded 15 °C (Figure S8). In females relative to males, genes involved in fatty acid transport (*cpt1a*) and anaerobic metabolism (*ldha*) were upregulated at 9 °C and 18 °C, respectively.

## Discussion

The extent of suitable thermal habitat for many fishes, including bull trout, is projected to decrease with global climate change (Sunday et al. 2012; Mochnacz et al. 2023), pushing many populations to their physiological limits. High-throughput transcriptional profiling is a sensitive and rapid approach to assess sublethal stress in fish, with machine learning classifiers facilitating interpretation of biomarker panels. Here, we show that mucus transcriptional profiling offers a minimally invasive approach to identify fish experiencing thermal stress. Transcriptional profiles in mucus were distinct between fish that experienced temperatures under 15 °C from those that were exposed to 15 °C and higher, which aligned with results in gill, liver, and muscle in juvenile bull trout. We identified the strongest combination of 10 tissue-specific gene biomarkers to inform a random forest classifier that resulted in over 90% classification accuracy of thermally stressed fish. Sex-biased gene expression was less pronounced in mucus and gill than in liver and muscle. Given its high predictive accuracy, we propose that mucus transcriptional profiling is a viable method to assess sublethal stress in wild fish and will be particularly important for species where lethal sampling is prohibited or when a less invasive approach is preferred by regulatory, research or community groups.

### Identifying thermally stressed fish using transcriptional profiling

The transcriptional thermal threshold in epidermal mucus reflected threshold estimates reported in gill, liver, and muscle in juvenile bull trout from the present study and of young-of-year acclimated to similar temperatures (Best et al. 2025). For each tissue in the present study, fish acclimated to 9 and 12 °C clustered separately from 15 and 18 °C, suggesting that shifts in transcriptional regulation in mucus can be leveraged to detect thermal stress. Interestingly, bull trout are cold-water specialists that generally occupy temperatures under 12 °C in the wild (Isaak et al. 2015; Mochnacz et al. 2023), which aligns with the cellular-level thermal thresholds reported thus far. Transcriptional thermal thresholds were also consistent with organismal-level thermal sensitivities relative to other species (Hinch et al. 2021), as the onset of cellular thermal stress occurred at a lower temperature in juvenile bull trout (15 °C) than sub-Arctic juvenile coho (21 °C) and Chinook salmon (23 °C) acclimated to 13 °C (von Biela et al. 2023).

Each tissue-specific biomarker panel resulted in high classification accuracy of bull trout according to thermal stress status. In all tissues, most genes in the reduced panel were associated with cellular stress activation when water temperatures exceeded 12 °C. This involved upregulation of molecular chaperones and genes involved in energy mobilization and apoptosis (de Nadal et al. 2011; Logan and Buckley 2015). Downregulation of anabolic pathways in favour of catabolic pathways, as well as decreased growth signaling and tissue remodelling in gill, liver, and muscle were further diagnostic of thermally stressed bull trout. Metallothionein b (*mtb*) emerged as a top biomarker of thermal stress across all tissues except gill in the present study. Upregulation of *mtb* in liver and muscle suggests a protective response against oxidative stress (Zeng et al. 2025), while its downregulation in mucus alongside upregulation of interleukin-8 (*il-8*) may promote a pro-inflammatory response (Gomez et al. 2013).

Three important biomarkers in mucus were involved in immune function, consistent with the role of mucus in host defences (Shephard 1994; Rombout et al. 2014). Mucus transcripts have been used to assess immune function in channel catfish (*Ictalurus punctatus*) following a bacterial challenge (Ren et al. 2015) and to detect immune suppression in juvenile mahi-mahi (*Coryphaena hippurus*) exposed to petroleum-derived chemicals (Greer et al. 2019), demonstrating that mucus transcripts are responsive to a myriad of stressors beyond warming temperatures. Furthermore, in temperature-acclimated lake trout (*Salvelinus namaycush*), transcripts for genes associated with immune function (*mhci*), stress responses (*mta*, *gstp1*), and osmoregulation (*atp1b1*) in mucus showed similar temperature-dependent patterns to the bull trout in our study (Weinrauch et al. 2025). This suggests that this suite of mucus biomarkers may be useful to detect thermal stress across different species, at least in salmonids. Similarly, heat shock protein 70 (*hsp70a*) transcripts in white muscle emerged as the top ranked biomarker for thermal stress in bull trout, consistent with findings in coho and Chinook salmon (von Biela et al. 2023). Taken together, these results provide support for developing pan-species biomarkers to assess sublethal stress using mucus transcripts, similar to gene panels developed for other tissues in salmonids (Akbarzadeh et al. 2024; Weinrauch et al. 2025; Banousse et al. 2025; Best et al. 2025; Islam et al. 2026).

### Sex-specific effects of thermal stress on transcriptional responses

Thermal stress responses were sex-dependent in the present study. In mucus and gill, responses to thermal stress were generally shared across sexes. However, subtle differences were detected in females relative to males, including greater thermal sensitivities in mucosal immune (*mhci*, *il8*) and stress responses (*mtb*). Differences in transcriptional responses to stressors in mucus due to sex are poorly understood and are likely life stage-, stressor-, and/or species-dependent. For example, interferon-related genes and antimicrobial peptide transcripts in epidermal mucus of juvenile yellowtail kingfish (*Seriola lalandi*) were less abundant in females than in males exposed to pathogens (Valero et al. 2024), whereas there was no distinct clustering of mucus transcriptional profiles by sex in lake trout exposed to diluted bitumen (Andrzejczyk et al. 2022). In gill, we detected earlier shifts in genes involved in glycolytic (*aldoaa*) and remodelling (*mmp2*) pathways in females relative to males, but overall transcriptional responses to thermal stress were similar between the sexes. Similarly, sex did not greatly influence the gill transcriptome of thermally stressed wild-caught adult pink (*Oncorhynchus gorbuscha*) and sockeye salmon (Jeffries et al. 2014). This suggesting that sex may modulate the magnitude of the gill transcriptional response, yet the pathways activated in response to thermal stress are conserved. The subtle sex differences in mucus and gill may enable classification with high accuracy without including sex to further inform models when using a non-lethal sampling design. This is important because sex determination of juvenile salmonids and younger adults by morphology alone can be difficult. For species with low sexual dimorphism like bull trout, this would require genetic sex determination using fin clip DNA to validate observations in the field (Amish et al. 2022).

In contrast, sex bias in cellular responses to thermal stress was more pronounced in liver and muscle, which may be related to sexual dimorphism in energy metabolism during periods of stress (Qiao et al. 2016). The liver is the primary site for energy substrate mobilization to support increased energy demands in peripheral tissues, such as gill and muscle, during stress (Antomagesh and Vijayan 2025). Accordingly, many temperature-responsive genes in liver were associated with growth and metabolism, consistent with findings in Atlantic salmon exposed to acute heat stress (Shi et al. 2019) and lake trout experiencing chronic heat stress (Weinrauch et al. 2025). Transcriptional shifts in liver occurred at lower temperatures in females relative to males, including downregulation of genes involved in lipid synthesis (*fasn*) and detoxification (*gstp1*) and altered expression of growth-related genes (*ghr*, *igf1*, *igfbp1*), indicative of earlier changes in metabolism in response to thermal stress. Conversely, transcriptional responses to thermal stress in muscle were delayed in females relative to males, where males showed earlier and sometimes greater upregulation of a suite of genes associated with the cellular stress response. Indeed, heat shock proteins and genes involved in apoptosis (e.g., *chmp5ab*) were upregulated at lower temperatures in the muscle of males, suggesting lower temperature sensitivity in their muscle relative to females. Taken together, transcriptional responses were tissue-specific but largely conserved between sexes in mucus and gill, whereas liver and muscle had more pronounced sex-specific thermal thresholds. Our results suggest that each sex may prioritize different metabolic tissues (i.e. liver vs. muscle) when coping with thermal stress, particularly during gonad development (Miller et al. 2009; Morash et al. 2013). Females in our study, had begun vitellogenesis which was associated with the consistent upregulation of liver carnitine palmitoyltransferase 1A (*cpt1a*) across all temperatures to fuel enhanced energy demands. It is worth noting that sex bias in gene expression in juvenile fishes is relatively understudied compared to adults but has been reported in as early as the embryo stage in rainbow trout (Hale et al. 2011).

## Conclusion

Epidermal mucus emerged as a viable matrix to identify thermally stressed individuals, as random forest classifiers trained with mucus transcriptional profiles performed comparably to classifiers using gill, liver, and muscle. Indeed, mucus represented transcriptional profiles intermediate to all tissues, bridging the distinct transcriptional profiles of gill, liver, and muscle. The positioning of mucus profiles closer to gill using multivariate approaches was also seen in lake trout (Weinrauch et al. 2025). This is likely due to both tissues interfacing directly with the aquatic environment and shared immune and osmoregulatory functions. Consideration of study objectives, however, will still be important when selecting tissues for transcriptional profiling. For instance, transcription of genes associated with growth and metabolism in mucus did not appear to be altered by thermal stress, suggesting that mucus may have limited use to answer questions about energy substrate metabolism to meet the enhanced energy demands of stress.

Transcriptional profiling of mucus to assess sublethal stress will be particularly important in species where minimally invasive approaches are preferred. Importantly, in the present study, mucus sampling did not lead to mortalities after 96 h. Collecting mucus samples without anaesthesia in wild populations would be comparable to the level of disturbance involved in routine measurements of fork length during fisheries surveys, which typically involves exposing fish to air for 30–60 seconds. As was demonstrated in juvenile gilthead seabream (*Sparus aurata*), mucus sample collection can be completed in under one minute with little effect on skin integrity and its antibacterial capacity (Sanahuja et al. 2023). Epidermal mucus is ideal to monitor sensitive species as it is continuously secreted, samples are collected rapidly, and it less invasive relative to dissections. Taken together, transcriptional profiling of epidermal mucus can be used to classify fish according to stress status, which is particularly important when monitoring sensitive species. Additionally, swabbing epidermal mucus requires little to no specialized training, which may facilitate enhanced sampling efforts and collaboration among diverse groups. Although non-lethal sampling approaches have been developed for many tissues (Jeffries et al. 2021), we propose that mucus is a promising matrix when the study objective is to obtain a systemic assessment of thermal stress. Our work provides a comprehensive comparison of transcriptional profiles in tissues that are commonly used to identify thermal stress in fishes and highlights the potential utility of mucus for transcriptional profiling for conservation-based research questions.

## Acknowledgements

We thank Kerri Wautier for their help in fish holding. We also thank Andrew Chapelski, Raven Bennett, Dr. Erik Folkerts, and Andrea Kneale for their help with tissue collection. Many thanks to Elli Hung for their assistance with sample preparation.

## Competing Interests

The authors declare there are no competing interests.

## Funding

This works was funded by a Mitacs Elevate Postdoctoral Fellowship to ALC (IT40703), a Fisheries and Oceans Canada Species at Risk Program project grant to NJM, and a Genome Canada Large-Scale Applied Research Project grant to KMJ.

## Author contributions

ALC, TD, and KMJ designed the research. ALC and TD conducted the experiment. BK, NJM, and KMJ acquired funding. ALC analyzed data and wrote the original draft. All authors reviewed and edited the final submission.

## Benefit-sharing

Benefits from this research come from openly sharing our data and findings on a public database as described above.

## Notes

### Competing Interest Statement

The authors have declared no competing interest.

### Summary of Updates

We have updated the Funding Information and Acknowledgements sections.

