## Supplemental Table and Figures for "Mucus transcriptional profiling as a minimally invasive approach to identify thermal stress in a stenothermal salmonid"

Table S1. Probes, primers, and tissue-specific PCR efficiencies for 56 genes measured in mucus, gill, liver, and muscle of juvenile bull trout (*Salvelinus confluentus*) using OpenArray qPCR. See Islam et al. (2026) for details on probe and primer design.

| **Gene symbol** | **Gene name** | **Probe** | **Forward primer** | **Reverse primer** | **Mucus** | **Gill** | **Liver** | **Muscle** |
|---|---|---|---|---|---|---|---|---|
| *aldoaa* | aldolase A | TGTACGAGCGGTGTGC | CAACGGAGAGACCACCACTCA | ACGCCACTTAGCAAAGTCAGC | 2.0 | 2.3 | 2.0 | 2.1 |
| *ampka1* | AMP-activated protein kinase | CGCAACCACCACGAC | AAGTTTGAGTGCACCGAGGAG | GACATAATGCGGCGGTTGTC | 2.0 | 2.1 | 2.1 | 2.0 |
| *atp1b1* | sodium-potassium ATPase (NKA) | CAGGCCCTGCTGCT | GGATGCTTGGCTGGGATTT | CTCTGTCCTGGTAGGTGGGTTT | 2.0 | 2.0 | 2.3 | 2.0 |
| *cam* | calmodulin | CGCTCTTTGACAAGGA | GAGGAGCAGATTGCCGAGTT | CATGACAGTGCCCAGCTCTTT | 2.1 | 2.4 | 2.2 | 2.1 |
| *casp3ab* | caspase 3b | CTGGTGGGCAAGCC | GGCGACCGCTGCAAAA | ACGCCTGGATGAAGAAGAGTTT | 1.9 | 2.0 | 1.8 | 1.8 |
| *casp9* | caspase 9 | TCCCAGCTTTAATAGAG | GCCAGACAGTTGGTTCGAGAC | GGCTATGCTGCCCTTTCTCA | 2.1 | 2.1 | 1.9 | 1.9 |
| *cat* | catalase | CGTCACACAGGTGCGTA | TGGGCCGCTACAACAGTACTG | CTCGTTCAGCACCTTAGTGAAGAA | 1.9 | 1.9 | 2.0 | 1.9 |
| *chmp5ab* | charged multivesicular body protein 5 | AGGATCTCCAGGACCAG | CCAAAGAGATGAAGGCGGC | TTGGCGTCCTCCATCATGT | 2.0 | 2.1 | 1.9 | 1.9 |
| *cirbpa* | cold-inducible RNA binding protein | CATTGGAGGGAATGAA | CGGGAAGGTCTCGTGGATT | TGGTTCTGCCATCGACAGACT | 2.1 | 2.2 | 2.1 | 2.1 |
| *clock1a* | Clock circadian regulator a | CTGGAGCAGAGGAC | CGCAATGCAGCACCTGAA | ATGTTGGCCTCGATCATCCT | 2.0 | 2.0 | 2.0 | 2.0 |
| *cpt1a* | carnitine palmitoyl transferase I | TCAATGACCCGGATGTT | TACAGCTGGCCCAATTCAGG | AACCGTCTCTGTCCTACCCTCA | NA | 2.0 | 2.1 | 2.2 |
| *cry1ab* | Cryptochrome circadian regulator 1a and 1b | TGGGCCGGTTAGC | CCGCCGGGACAAGGA | AGATGATTTCTACTCCGTGCTCTTC | 1.8 | 1.9 | 1.9 | 1.9 |
| *cs* | citrate synthase | CAGCAGCATTGGC | TTGATTGCCAAGTTGCCGT | ATGTTGGCGAAGTTAGCGGA | 2.0 | 2.0 | 2.0 | 2.0 |
| *ctsd* | cathepsin D | CCACAAGTATAACGGTGCC | TTCACAGACATCGCCTGCTT | GGTACCCAGACAGACTGCCAG | 1.9 | 2.1 | 1.9 | 1.9 |
| *cyp1a* | cytochrome P450 1a | CCCGGAGCTGTGGAA | CAGTGGCAGGTCAACCATGA | CAGCACTCAGGAAACGGTCA | 1.9 | 2.0 | 2.1 | 2.0 |
| *ef1a* | elongation factor alpha | CCGCCATCATCGTCAT | GAAGCTTGAGGACAACCCCA | GAAGCTCTCCACACACATGGG | 2.0 | 2.0 | 2.1 | 2.3 |
| *fasn* | fatty acid synthase | AGGTGGAGGAGGCC | TGTGGGAGGTGTAGTCAAGCC | TCCCTGGGCCATGTATCTGA | 1.9 | 1.9 | 2.0 | 1.9 |
| *ghr* | growth hormone receptor | TGAGCTGGGAGCCG | TAACCGGGAGCCACTTTGAC | ATTGACCTCACGGTACTGCACC | 2.1 | 2.0 | 2.0 | 2.0 |
| *gr2* | glucocorticoid receptor 2 | AGACTGCAGGTGTCTCA | ATGGCAGACCAGTGTGAACAGAT | GAGCAGCAGCAGAACCTTCAT | 1.9 | 1.8 | 1.9 | 1.8 |
| *gstp1* | glutathione-s-transferase pi isoform | TGAAGTGCTGCTCAAC | GGTGACAAGCCTTCGTTTGC | CAAAGCTCTTCAGGGAGGGG | 2.1 | 2.2 | 2.2 | 2.0 |
| *hk1* | hexokinase 1 | TCCTCTCCCAGATTGA | CCAGAGGCATTTTCGAAACTAAG | AGCGCCAACCGGTCACT | 1.9 | 2.0 | NA | NA |
| *hsd11b2* | 11β-hydroxysteroid dehydrogenase type 2 | CCTGTTCATCAACACACT | CTGTCTAGCAGCGTACGGAGC | ATGGTGGACACTTTGACCCC | NA | 2.0 | 2.0 | 2.1 |
| *hsf1* | heat shock factor 1 | TGCAGGGCTCGCCT | CCCAAGTTCAGCAGGCAGTAC | GCCGTGAAGAGACCGGTACT | 1.9 | 2.1 | 2.0 | 2.0 |
| *hsp70a* | heat shock protein 70a | ACACCTCCATCACCAGG | GAGAACACTGTCCTCCAGCTCC | CCCTGAAGAGGTCGGAACAC | 1.9 | 2.1 | 2.0 | 2.1 |
| *hsp7c* | heat shock cognate 71 kDa protein | ACCCAGTTATGTCGCCTT | TAGCCAACGACCAGGGAAAC | CGTCCCCAATCAGCCTTTCT | 2.0 | 2.1 | 2.1 | 2.0 |

Table S1. (*continued*)

| **Gene symbol** | **Gene name** | **Probe** | **Forward primer** | **Reverse primer** | **Mucus** | **Gill** | **Liver** | **Muscle** |
|---|---|---|---|---|---|---|---|---|
| *hsp90ba* | heat shock protein 90 (constitutive) | TCTCCAGGGACACAGAC | GAGGTGGAGGAGGACGAGTACA | GCTGTGAAGTGGATGTGGGA | 2.0 | 2.1 | 1.9 | 1.8 |
| *hspa4* | heat shock 70 kDa protein 4 | ACCTGTGGCTGATTGT | ACTGCTGAGACCGCAATGAA | TGCGTCAGTGTAGAAGCTGGG | 2.1 | 2.1 | 2.0 | 2.1 |
| *igf1* | insulin-like growth factor 1 | ACACCCTCTCACTGCT | TTCAAGAGTGCGATGTGCTGT | CGCCGAAGTCAGGGTTAGG | NA | 2.0 | 2.2 | 2.1 |
| *igf2* | insulin-like growth factor 2 | TGCAGTTCGTCTGTGAAG | ATGTGGAGGAGAACTGGTGGAC | CCTGCTGGTTGGCCTACTGA | 1.9 | 2.0 | 2.1 | 1.9 |
| *igfbp1* | igf-binding protein 1 | TGGACAGAGAGGCAGGT | CCAAACAGTGTGAGTCGTCTCTTG | TCCAGGAGGACACACACCAA | NA | 1.8 | 1.9 | 1.9 |
| *il1b* | interleukin-1 beta | ACCCCATCACCATGC | TCAGGGTCTGGATCTGGAGG | CTCGGTTCCCATGGTAACCC | 1.1 | 1.7 | NA | NA |
| *il8* | interleukin-8 | ATGAGTCTGAGAGGCATG | TGGCCCTCCTGACCATTACT | GTCTCAATGCAGCGACATCG | 2.0 | 2.0 | NA | NA |
| *ldha* | lactate dehydrogenase a | TGCTGGTCGTCTCCAA | CGTCAAGTACAGCCCCAACG | GCCACGTAGGTCAGGATGTCA | NA | NA | NA | 2.1 |
| *ldhb* | lactate dehydrogenase b | CCAACCCAGTGGACGT | CCCCCAACTGCACCCTTATT | AATCCGCTCAACTTCCACGT | 2.0 | 2.1 | 2.0 | 2.0 |
| *lepr* | leptin receptor | CCGGGACCTGGAGTGA | CGCTGTATGCACATCAACGG | GTCAGGAGCTCTGCTGTTGTGA | NA | 2.0 | NA | 2.0 |
| *lipeab* | hormone-sensitive lipase a/b | CTTCTGCACATCATCCA | ACACTGGTCAAGGTGTTGCAGT | TGCAGTTAGCGGCAATGTAGC | 2.2 | 2.1 | 2.1 | 2.2 |
| *lpl* | lipoprotein lipase | TGGACTGGCTGACACGG | TGACAGCGCTGTACAAGAGGG | GGAGGTGAGGTAGTGCTGCTG | NA | 2.0 | 2.2 | 2.1 |
| *mhci* | major histocompatibility complex class 1 | AGTGACCTGCCACGCG | AGTCCCTCCCTCAGTGTCTCTG | AGGACACCATGACTCCACTGG | 2.2 | 2.3 | 2.2 | 2.1 |
| *mmp2* | matrix metalloproteinase 2 | AGGAAACCCAAGTGGCA | CGCTGTGGAGTTCCTGATGTT | AGGTCAGGAGAGTGGCCTAGAA | NA | 2.2 | 2.1 | 2.1 |
| *mmp9* | matrix metalloproteinase 9 | CACCTCCAAGGCTC | GCAGCGGTTTCCAGTCCAT | CATCTGCCTCTGCATTCTCATC | 1.6 | 2.1 | 2.0 | 1.4 |
| *mr* | mineralocorticoid receptor | CCAGCAGCAACACA | CATAGTCAATGTCAGCTGCTCCC | GCTGCTGCTCTGGCTTCTTCT | 2.0 | 1.9 | 2.0 | 2.0 |
| *mta* | metallothionein a | CGGTGGATCCTGCAAG | TGGATCCTTGTGAATGCTCCA | CTTACAACTGGTGCATGCGC | 2.1 | 2.1 | 2.2 | 2.1 |
| *mtb* | metallothionein b | TTCAGGCTGTGTGTGCA | GAAAAGTTGCTGCCCCTGC | AACAGCTGGTATCGCAGGTCTT | 2.2 | 2.2 | 2.3 | 2.1 |
| *nfe2l2a* | nuclear factor erythroid 2 like 2 | AGTAACCCTGGGAGTGC | GGTGGCTACAGCGATTCAGAC | GAAAGAGGTGTCTGCAGGCC | 2.0 | 2.1 | 2.0 | 2.0 |
| *pck1* | Phosphoenolpyruvate carboxykinase | CGTGTTTGTCGGAGCT | AGGCCTTTAGCTGGCAACAC | AGCTGTGGCTTCTGATCTCATG | NA | NA | 2.0 | NA |
| *pdcd10ab* | programmed cell death protein 10 | TGACCCAGGACATCAT | AGGCTGAGAAGGAGAACCCAG | CACATCGTCTGCAGCCATTC | 2.0 | 2.1 | 2.0 | 2.1 |
| *pgk* | phosphoglycerate kinase | CCTTCACCTTCCTCAATG | AGATGATCATCGGTGGTGGC | CATACAGGGAGGTGCCGATC | 2.0 | 2.2 | 2.1 | 2.0 |
| *rpl13a* | ribosomal protein L13a | CATGGTCGTACCTGCT | CACTGGAGAGGCTGAAGGTGT | GTGGGCTTCAGACGGACAAT | 2.1 | 2.1 | 2.2 | 2.0 |
| *rpl7* | 60S ribosomal protein L7 | CAAGGTGCGCAAGGT | TCGTCATCAGGATCAGGGGT | GAAGATCTGACGCAGACGCA | 2.0 | 2.1 | 2.1 | 2.0 |
| *rps9* | 40S ribosomal protein S9 | CCCGGCCGTGTCAA | TTCTCCCTGCGTTCACCATAC | GGCCCTTCTTGGCATTCTTT | 2.0 | 2.1 | 2.1 | 2.0 |
| *serpinh1* | serine proteinase inhibitor H1 | CGGCCAAGTCCACCGA | AGCGCTGTGAAGTCCATCAA | TGATGATCATGGCCCCATC | NA | 2.0 | 2.0 | 2.0 |

Table S1. (*continued*)

| **Gene symbol** | **Gene name** | **Probe** | **Forward primer** | **Reverse primer** | **Mucus** | **Gill** | **Liver** | **Muscle** |
|---|---|---|---|---|---|---|---|---|
| *slc2a1a* | solute carrier family 2 member 1a | TTGGCCCGGGTCC | TGAGCATCGTGGCCATCTT | AACAGCTCAGCCACGATGAA | 1.9 | 2.0 | 1.7 | 1.9 |
| *sod1* | superoxide dismutase 1 (Cu-Zn type) | CGGACCCCACTTCA | CAACACCAACGGCTGTATGAGT | CCTCCGTGGGTCTTGTTGTG | 2.0 | 2.1 | 2.1 | 2.1 |
| *sod2* | superoxide dismutase 2 (Mn-type) | AAGCTCCGTATCACAGC | GATGGCTGGGCTTTGACAA | CCTGCAGTGGGTCTTGATTAGG | 1.7 | 1.7 | 2.0 | 2.1 |
| *stat1* | signal transducer and activator of transcription 1 | CGGGCCCTGTCACT | CAGAAAGGCTTCCTGGAGGG | CTCTGTGGATGTGTGGGCAT | 1.9 | 2.1 | 2.0 | 2.0 |
| *vegfc* | vascular endothelial growth factor c | TGCTTCCTGAAAGGCAA | TGTGCCTGCGAGTGCAA | GGTGGCCTGGTGGAACCT | NA | 1.8 | 1.7 | NA |

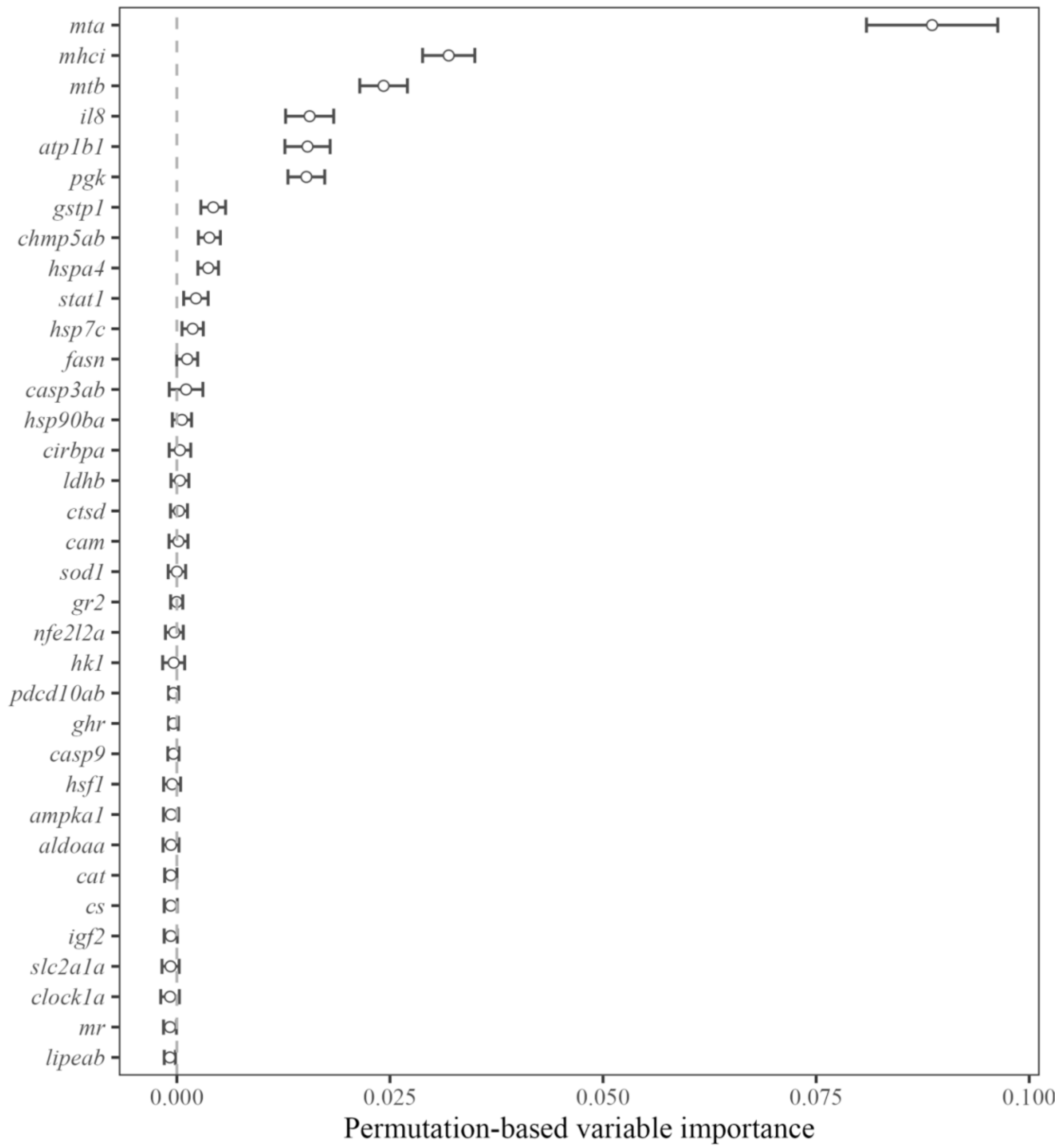


Figure S1. Mucus gene biomarkers ranked by permutation-based mean decrease in accuracy from random forest classifier used to distinguish bull trout (*Salvelinus confluentus*) thermal stress status. Points show means and bars indicate standard deviation across 1000 trees.

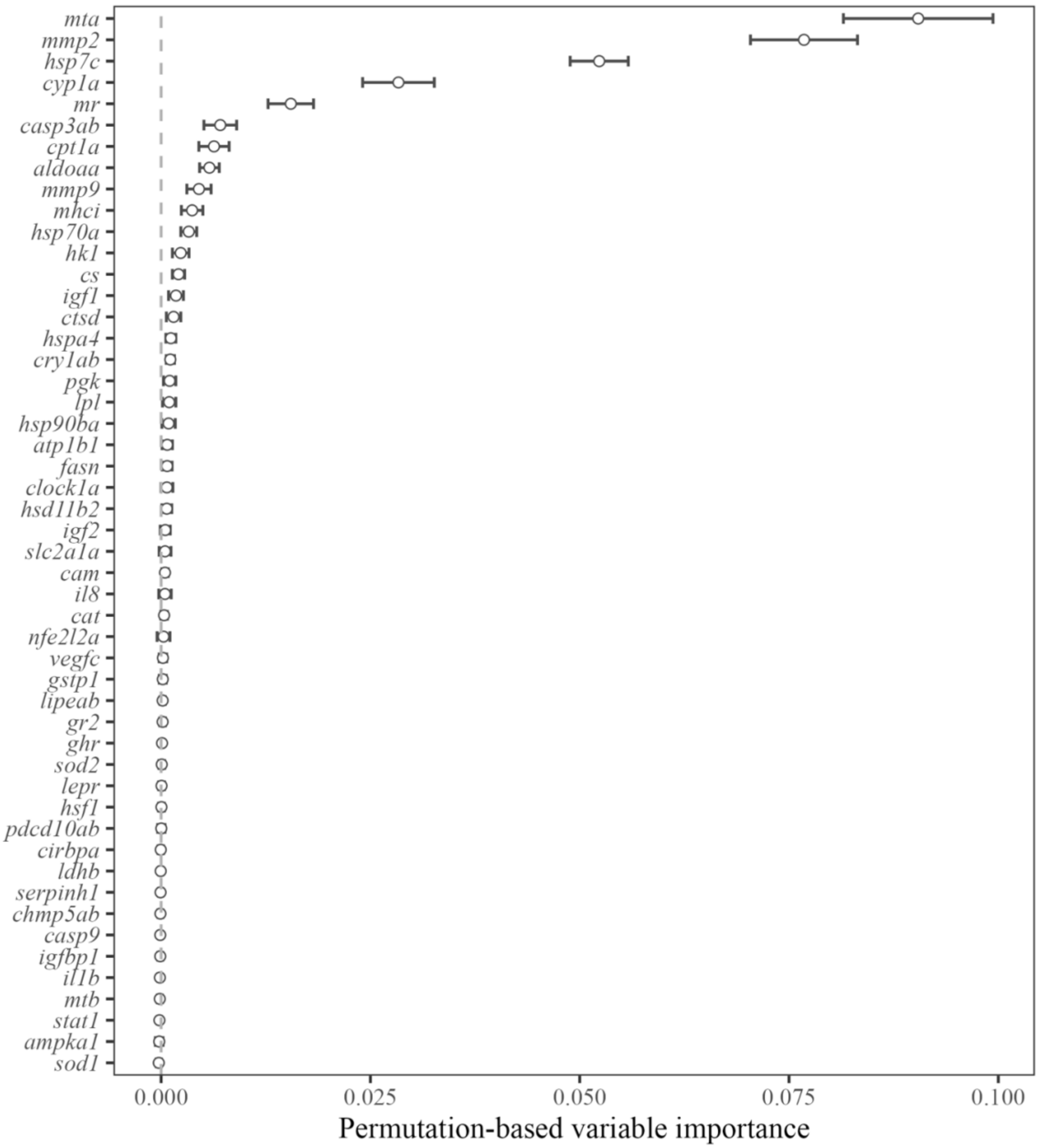


Figure S2. Gill gene biomarkers ranked by permutation-based mean decrease in accuracy from random forest classifier used to distinguish bull trout (*Salvelinus confluentus*) thermal stress status. Points show means and bars indicate standard deviation across 1000 trees.

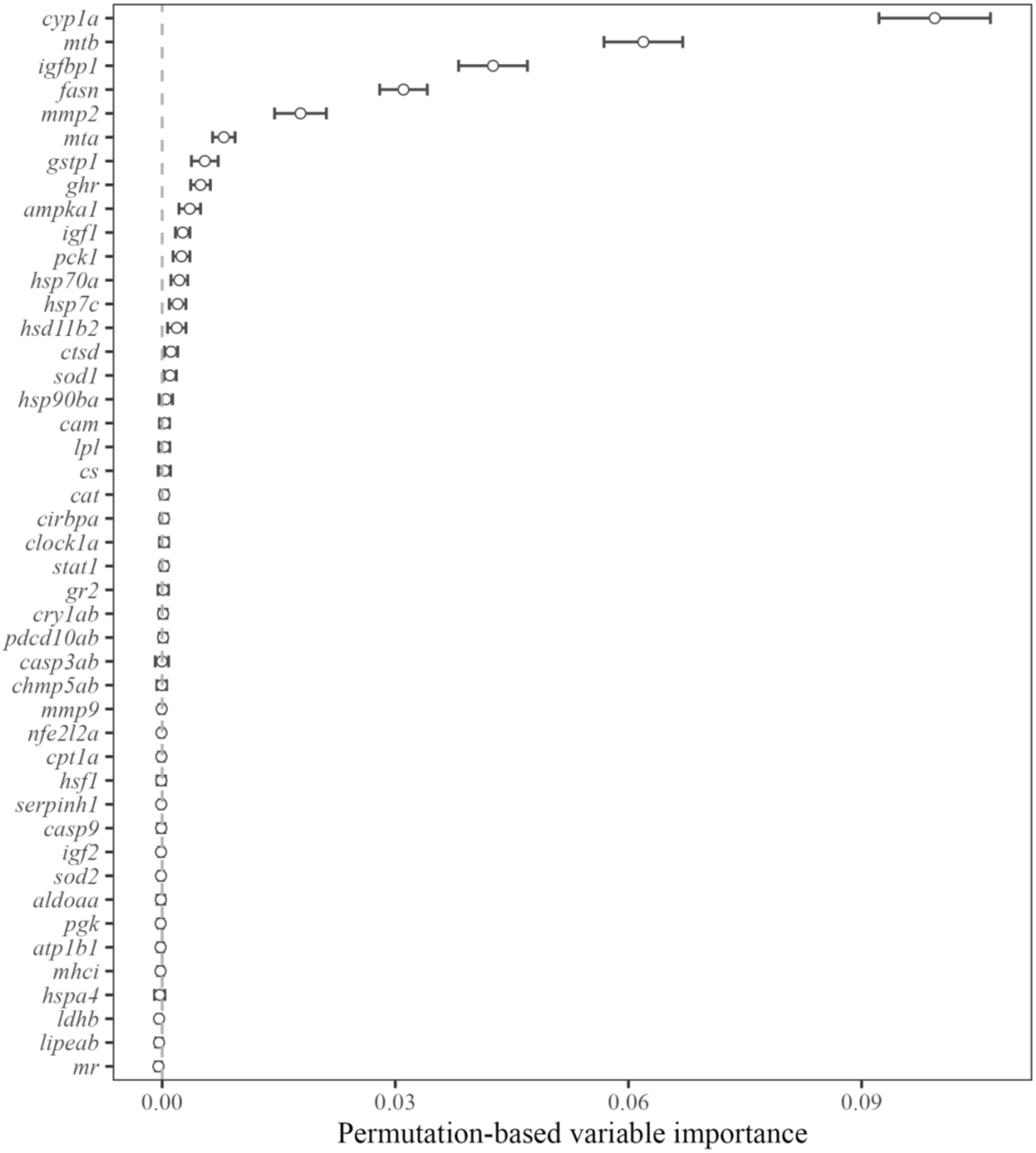


Figure S3. Liver gene biomarkers ranked by permutation-based mean decrease in accuracy from random forest classifier used to distinguish bull trout (*Salvelinus confluentus*) thermal stress status. Points show means and bars indicate standard deviation across 1000 trees.

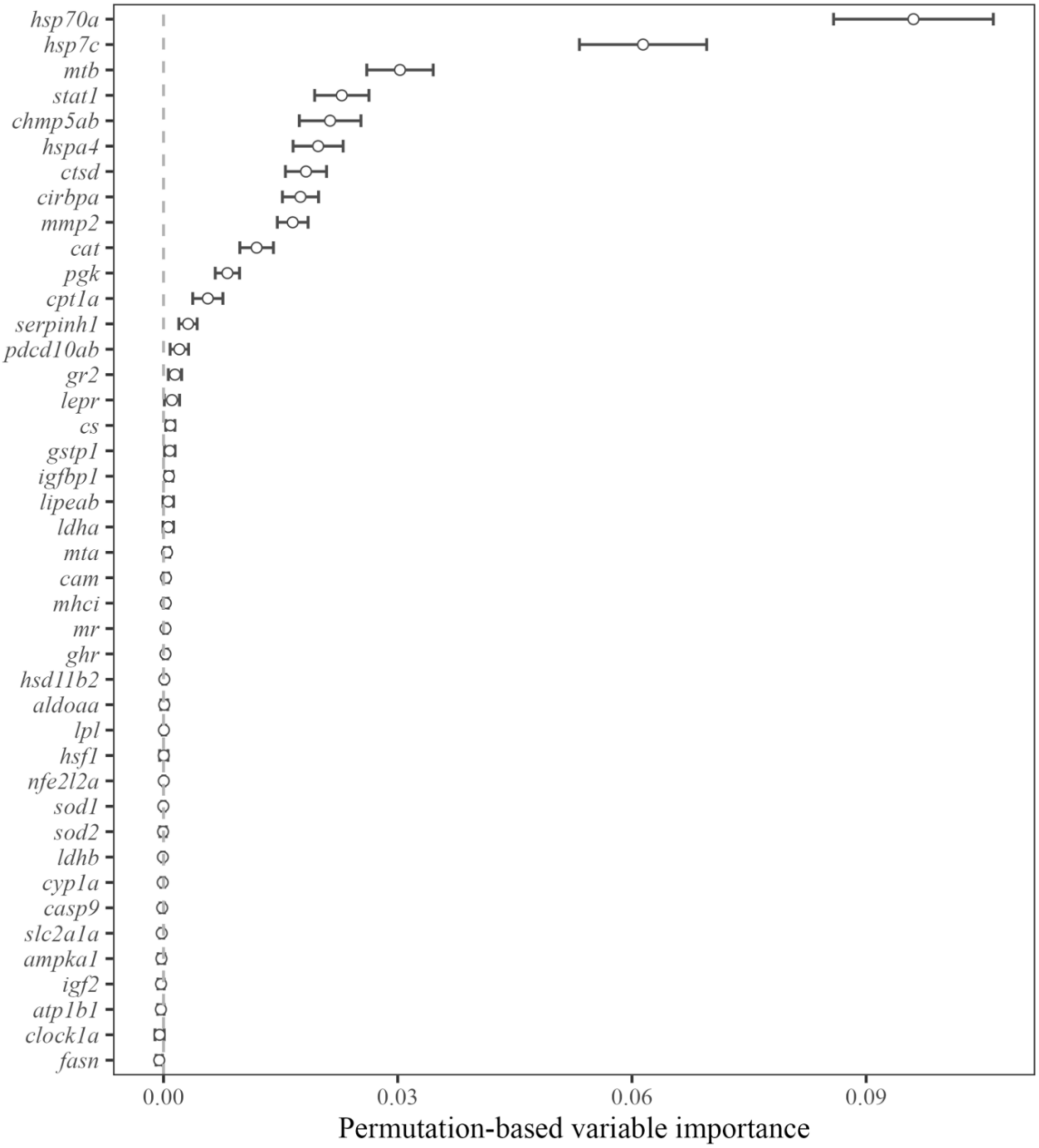


Figure S4. Muscle gene biomarkers ranked by permutation-based mean decrease in accuracy from random forest classifier used to distinguish bull trout (*Salvelinus confluentus*) thermal stress status. Points show means and bars indicate standard deviation across 1000 trees.

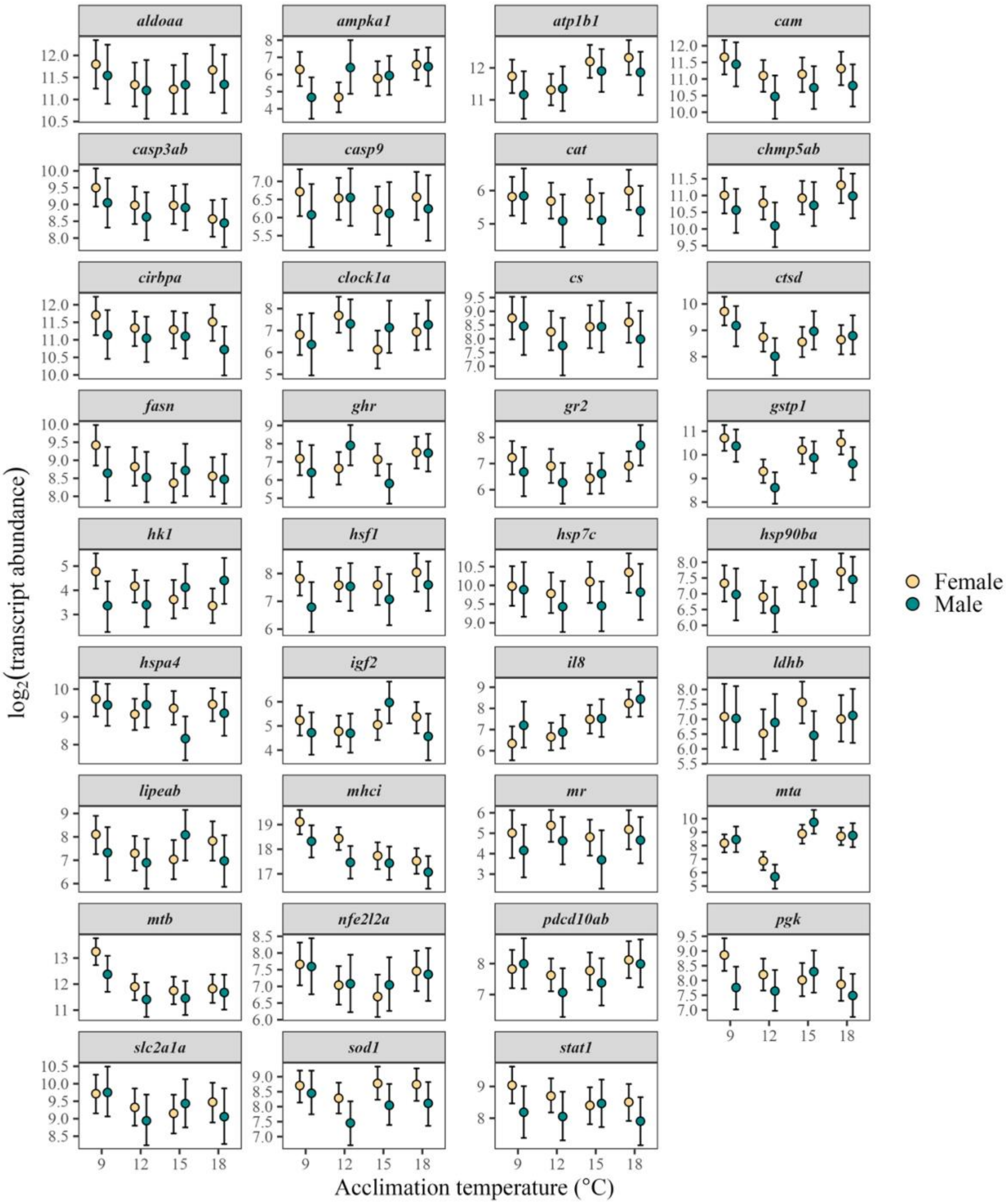


Figure S5. Sex-specific mucus transcriptional response of 35 genes in bull trout (*Salvelinus confluentus*) acclimated to 9, 12, 15, or 18 °C for four weeks. A single Bayesian generalized linear mixed model was used to analyze transcript abundance in mucus. Points represent posterior means and bars show 95% credible intervals.

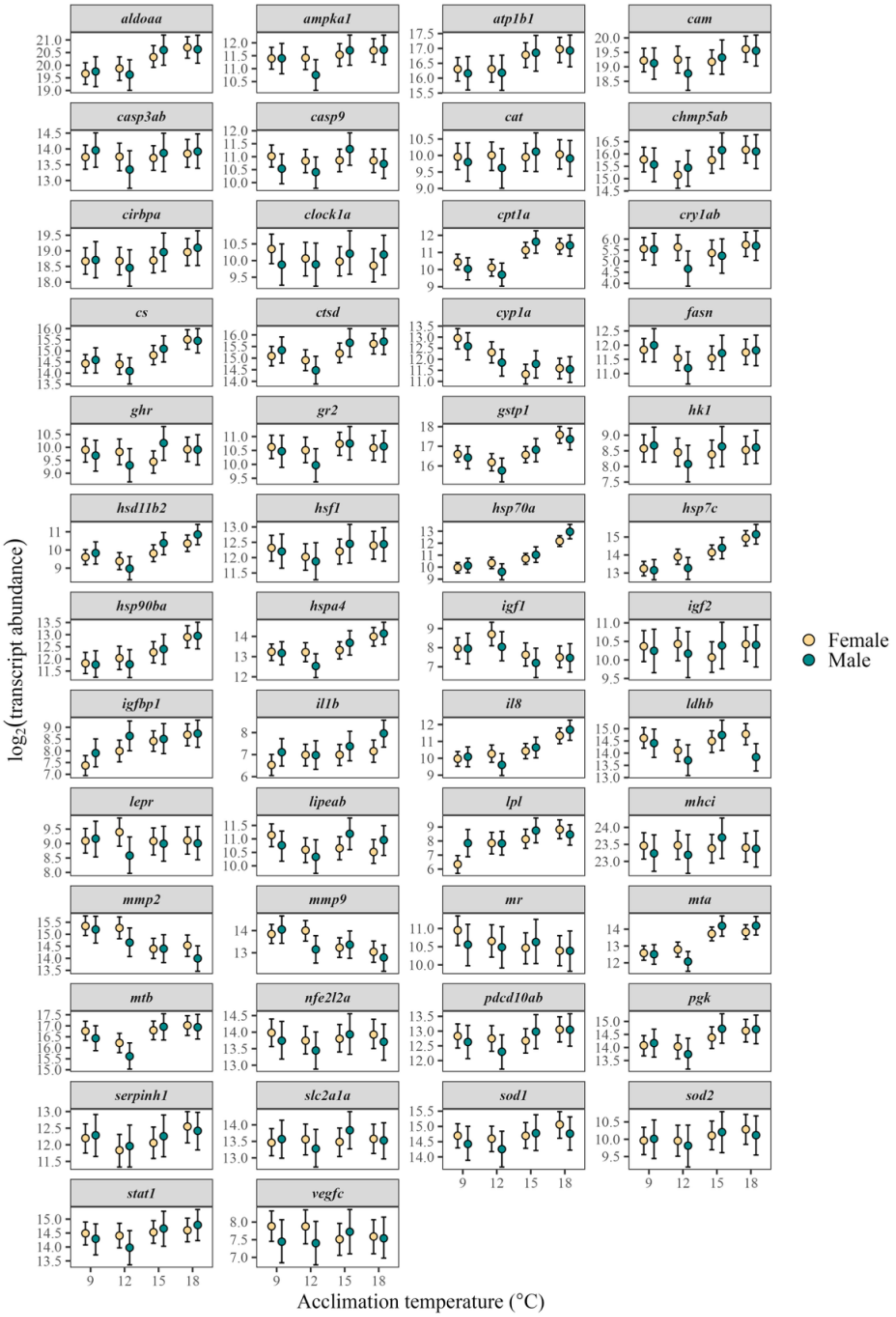


Figure S6. Sex-specific gill transcriptional response of 50 genes in bull trout (*Salvelinus confluentus*) acclimated to 9, 12, 15, or 18 °C for four weeks. A single Bayesian generalized linear mixed model was used to analyze transcript abundance in gill. Points represent posterior means and bars show 95% credible intervals.

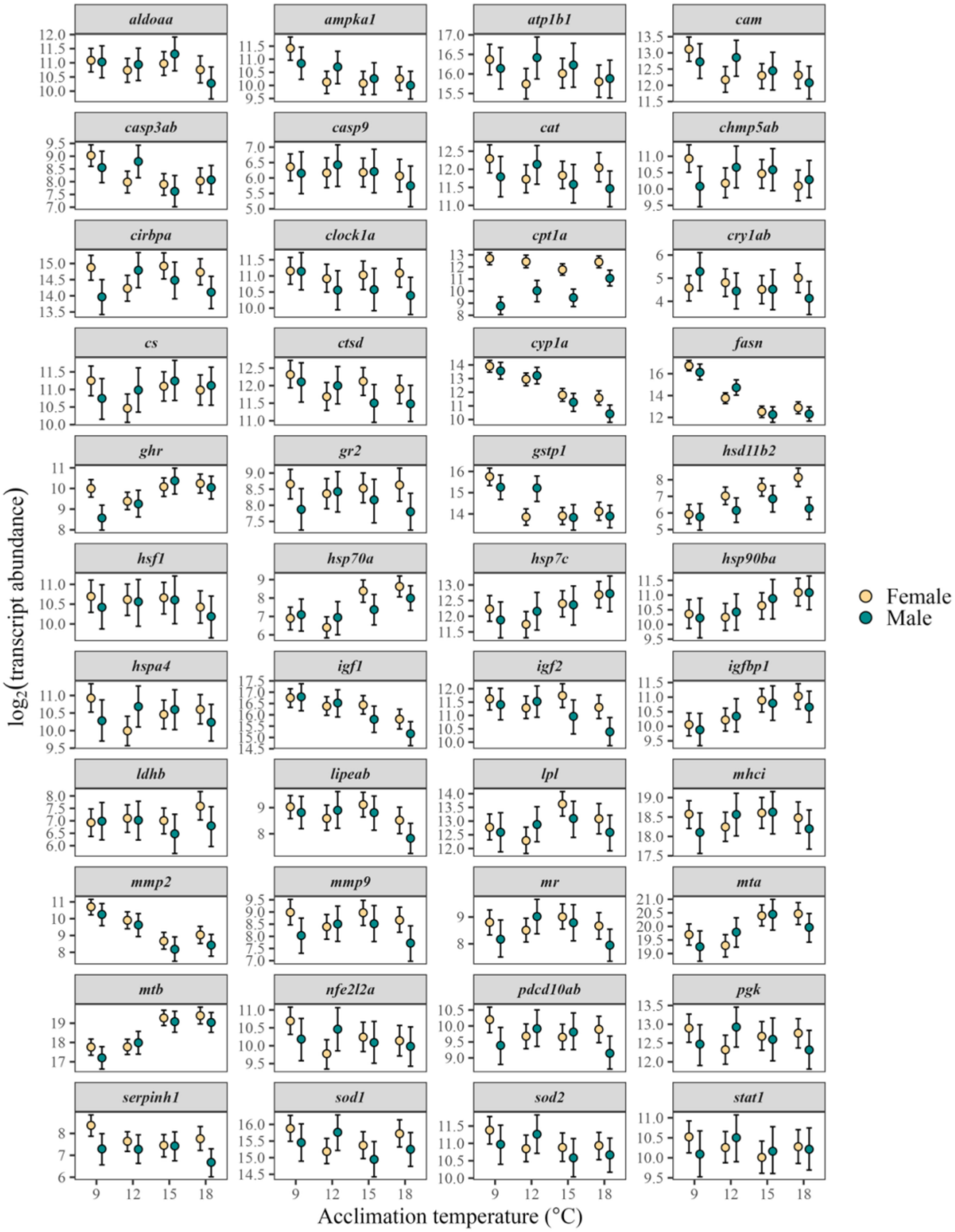


Figure S7. Sex-specific liver transcriptional response of 44 genes in bull trout (*Salvelinus confluentus*) acclimated to 9, 12, 15, or 18 °C for four weeks. A single Bayesian generalized linear mixed model was used to analyze transcript abundance in liver. Points represent posterior means and bars show 95% credible intervals.

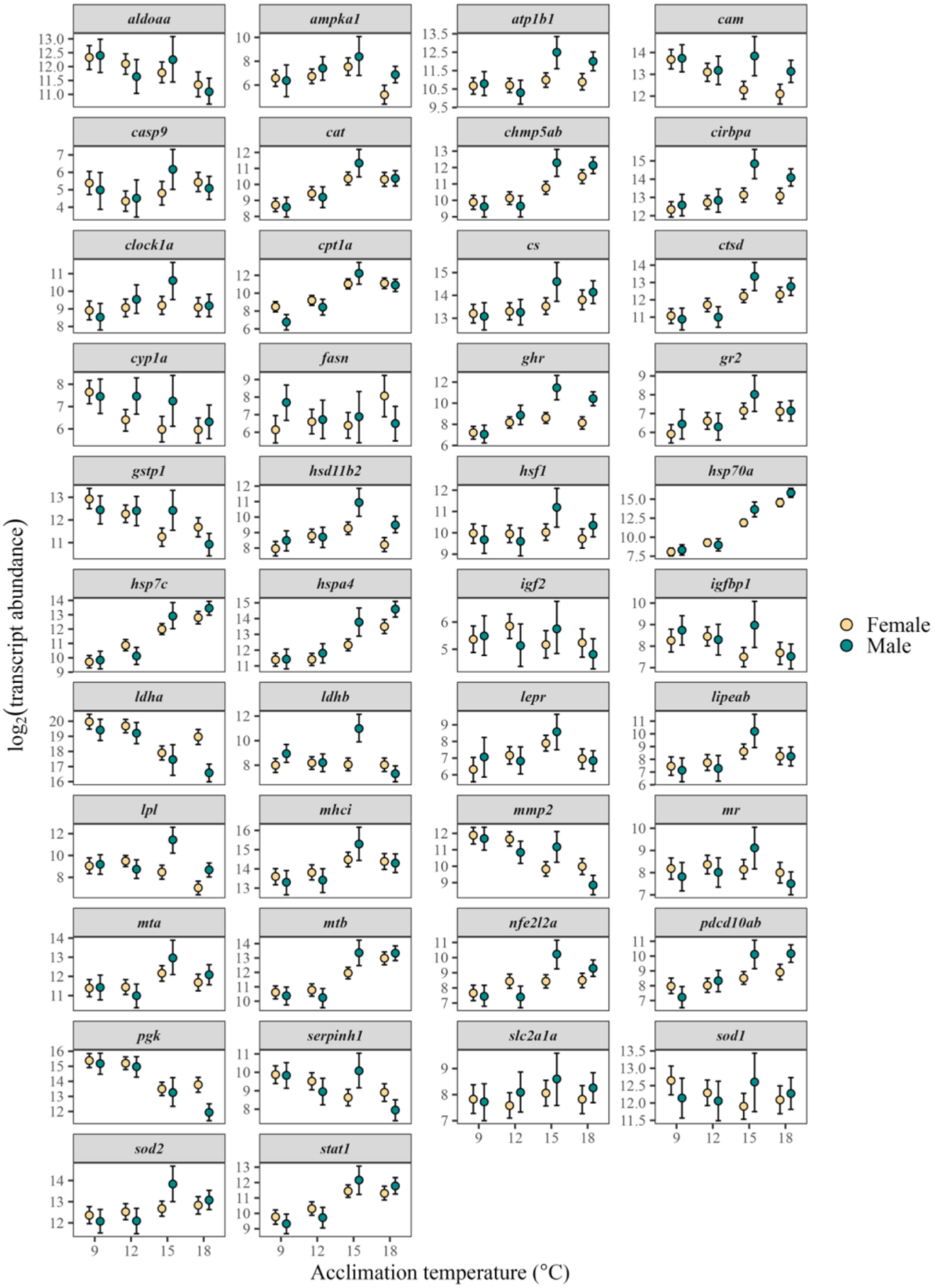


Figure S8. Sex-specific muscle transcriptional response of 42 genes in bull trout (*Salvelinus confluentus*) acclimated to 9, 12, 15, or 18 °C for four weeks. A single Bayesian generalized linear mixed model was used to analyze transcript abundance in muscle. Points represent posterior means and bars show 95 % credible intervals.
